# *C. elegans* behavior, fitness, and lifespan, are modulated by AWB/ASH-dependent death perception

**DOI:** 10.1101/2024.10.07.617097

**Authors:** Mirella A. Hernandez-Lima, Brian Seo, Nicholas D. Urban, Matthias C. Truttmann

## Abstract

The ability of the nervous system to initiate intricate goal-directed behaviors in response to environmental stimuli is essential for metazoan survival. In this study, we demonstrate that the nematode *Caenorhabditis elegans* perceives and reacts to dead conspecifics. The exposure to *C. elegans* corpses as well as corpse lysates activates sensory neurons AWB and ASH, triggering a glutamate-and acetylcholine-dependent signaling cascade that regulates both immediate (aversion) and long-term (survival) responses to the presence of a death signature. We identify increased adenosine monophosphate (AMP) and cysteine concentrations as chemical fingerprints for the presence of metazoan corpses and show that death cue sensing by AWB and ASH leads to physiological changes which promote reproduction at the expense of lifespan. Our findings illuminate a novel signaling paradigm that allows organisms to detect and interpret the environmental enrichment of intracellular metabolites as a death cue.

## Introduction

Metazoans employ conserved yet needs-adapted sensory systems to survey and interpret their immediate environments. These systems integrate information presented by both rudimentary (smell, pressure, temperature) and complex (e.g., the local abundance of conspecifics) sensory cues to facilitate an appropriate physiological and behavioral response. In this study, we were interested in defining the mechanistic and neuronal basis for the perception of dead conspecifics, or death perception, in the nematode *Caenorhabditis elegans*. An increasing number of species across the animal kingdom are recognized to perceive dead individuals of the same species, an event promoting a range of distinct responses. For example, social insects such as ants, honeybees, and termites recognize dead colony members and remove them from their hives or nests to maintain hygienic conditions and prevent the spread of infectious diseases within the colony^1–3^. This response is dependent on the release of oleic acid from the decaying corpses^4^. The vinegar fly, *Drosophila melanogaster,* recognizes dead conspecifics based on both visual and olfactory cues. These cues activate serotonergic signaling in ring neurons of living conspecifics, promoting aversive behaviors and decreased lifespan^5,6^. In zebrafish, the scent of dead conspecifics increases cortisol levels and induces defensive avoidance behavior^7^, whereas in Western scrub-jays (*Aphelocoma california*) the sight of a dead conspecific is sufficient to induce risk-reducing behavior and alarm calling^8^. Primates exhibit increased vocalization, aggressive displays, and, in most cases, an urge to inspect dead conspecifics without touching them^9,10^. In humans, the experience of encountering a deceased individual can promote complex, context-dependent physical (headaches, fatigue, nausea) and emotional (fear, anxiety, depression) responses^11,12^. These examples suggest that the perception of dead conspecifics may signal danger and lead to risk-mitigating and fear-induced behavioral outputs, affecting internal states^13^. The sensory modalities through which different species perceive dead conspecifics, and how this experience affects organismal health and fitness are poorly characterized. It is further unknown if there is an evolutionary conserved “universal death signature” that may facilitate the detection of corpses from different taxa.

In this study, we establish the physiological and mechanistic basis for a signaling paradigm that is triggered in response to the presence of metazoan corpses in *C. elegans*. We show for the first time that olfactory neurons AWB and ASH sense the environmental presence of corpses or corpse lysates. Downstream propagation of the sensory information promotes aversive behavior and increases local egg laying but shortens lifespan. We further demonstrate that this signaling paradigm requires neurotransmitters glutamate and acetylcholine, in parallel to GPCR- and adenylyl cyclase-mediated signaling transduction. Exposure of *C. elegans* to corpse lysates of distinct metazoan species decreases their health- and lifespan, suggesting that this response is both ancient and conserved. Finally, we identify the environmental presence of the nucleotide AMP, and the amino acid cysteine as potential death cues. Taken as a whole, our data provide novel insights into a novel sensory paradigm that modulates health, aging, and physiology in response to death perception.

## Results

### Exposure to dead *C. elegans* promotes aversive behavior in naïve individuals

Previous studies in *C. elegans* established that environmental cues, including crowding, the abundance of sex-specific pheromones, and the presence of attractants and repellants influence internal states, which affect behavior and health^14–19^. We thus hypothesized *C. elegans* could elicit a behavioral response when encountering dead conspecifics. To test this hypothesis, we performed choice assays in which we tested whether the presence of *C. elegans* corpses in an *E.coli* OP50 lawn would prevent adult worms (choosers) from feeding in these lawns. We devised an experimental setup in which 30 choosers were released in the center of an assay plate that contained four equally sized and distributed OP50 lawns in four quadrants (**Figure 1A, upper panel**). The distribution of choosers was scored three hours post release in the center of the plate. Control experiments confirmed that worms did not show a positional preference, distributing equally among the four bacteria lawns in the absence of any corpses (**Figure S1A**). The addition of approximately 50 naturally deceased worms to two of the four lawns induced a significant avoidance response towards OP50 lawns containing corpses (**Figure 1B**). We also obtained similar results when using corpses of synchronized day 1 adult worms that were killed by the exposure to sodium azide (NaN_3_) and vigorously washed to remove any NaN_3_ prior to placing them into the bacterial lawns (**Figure 1C**). The avoidance phenotype was further confirmed in classic avoidance assays, in which choosers showed an approximately 30%, reduction in bacteria lawn occupation in response to the presence of worm corpses (**Figure S1B-C**). The effect size was dose-dependent and proportional to the number of worm corpses present (**Figure S1D**). In a next step, we tested choosers in a four-option choice paradigm in which one OP50 lawn was spiked with M9 buffer, serving as a control, whereas the other three OP50 lawns were supplemented with 10μg of lysates from synchronized day-1 adult animals that were killed using sodium azide (CL1), lysed without prior inactivation (CL2), or killed in 100% ethanol before lysis (CL3). (**Figure 1A, lower panel**). We observed that choosers sorted non-randomly and significantly avoided feeding on bacterial lawns containing any corpse lysates (**Figure 1D**). Consistent with previous experiments (**Figure S1D**), we found that this avoidance phenotype was dose dependent, with at least 2.5μg of worm lysate per OP50 lawn required to trigger an aversive response (**Figure 1E**). Worm corpse lysate was equally potent in inducing an aversive response as the presence of *Pseudomonas aeruginosa* isolate PA14, a bacterial pathogen known to induce avoidance behavior in *C. elegans* (**Figure S1E**). This underscores the potency of worm corpse lysates to induce behavioral changes. Control experiments using chemically inactivated *E.coli* OP50 as food source further confirmed that components within the *C. elegans* corpse lysates directly promoted the observed behavior (**Figure 1F**). Taken as a whole, these results establish that the nematode *C. elegans* recognizes and reacts to the presence of dead conspecifics.

**Figure 1.**
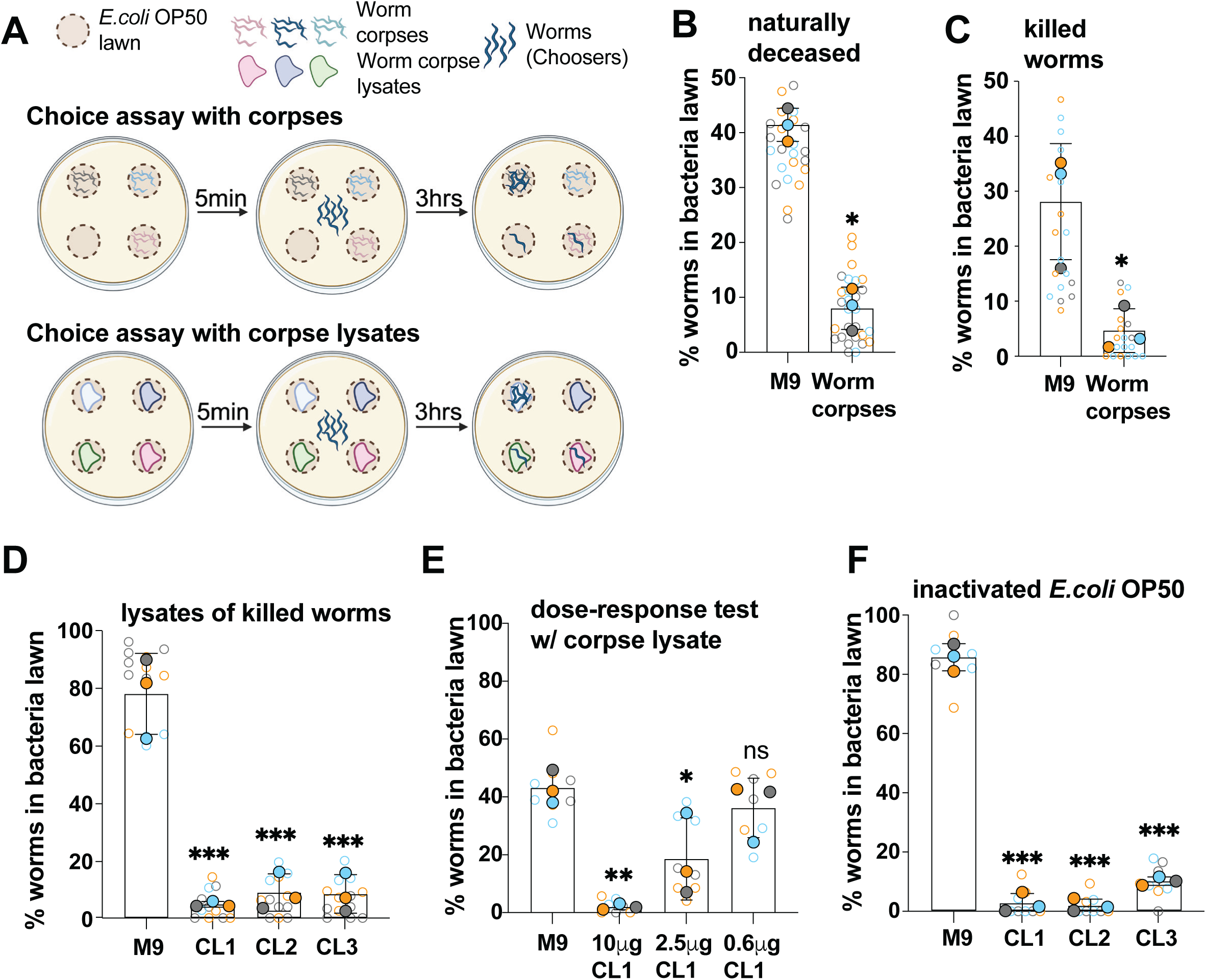
*C. elegans* recognize and avoid conspecific corpses. (A) Schematic of four-option choice assays used in this study. (B-C) Four-option choice assays testing worm behavior upon exposure to naturally-deceased (B) and chemically inactivated (C) worms. Corpses were added to two of the four *E.coli* OP50 lawns whereas the two remaining *E.coli* OP50 lawns were supplemented with M9 and served as controls. Corpse to chooser ratio was 2:1. (D) Four-option choice assays testing worm behavior upon exposure to worm corpse lysates. Three *E.coli* OP50 lawns were supplemented with 10μg worm lysate whereas the fourth *E.coli* OP50 lawn was supplemented with M9 and served as control. CL1: lysate of NaN_3_-inactivated worms; CL2: lysate of worms lysed without prior inactivation; CL3: lysate of ethanol-inactivated worms. (E) Four-option choice assays testing worm behavior upon exposure to different concentrations of worm corpse lysates of NaN_3_-inactivated worms. Three *E.coli* OP50 lawns were supplemented with indicated amounts of worm lysate whereas the fourth *E.coli* OP50 lawn was supplemented with M9 and served as control. (F) Four-option choice assays testing worm behavior upon exposure to worm corpse lysates. All four *E.coli* OP50 lawns were prepared using PFA-inactivated *E.coli* OP50 bacteria. Three *E.coli* OP50 lawns were supplemented with 10μg worm lysate whereas the fourth *E.coli* OP50 lawn was supplemented with M9 and served as control. For (B) through (F): Solid circles represent individual biological replicates; hollow circles represent individual technical replicates; corresponding biological and technical replicates are color matched. Error bars represent standard deviation of the mean. Indicated P values were calculated using unpaired two-sided t tests (B-C) and 1-way ANOVA tests with multiple comparison (D-F, with M9 control condition serving as reference). ns=p>0.05 (not significant); *p=<0.05; **p=<0.01; ***p=<0.001.

### Factors influencing aversive behavior

To better understand this new behavioral phenotype in *C. elegans*, we performed subsequent experiments in which we tested how parameters such as age of choosers, age of inactivated worms, feeding status, food source, and biological sex affect the observed choice to avoid dead conspecifics. Using the previously introduced corpse lysate-based choice paradigm, we found that neither the age of adult choosers (**Figure 2A, S2A-B**) nor the age of the worms used to prepare the lysates (**Figure 2B, S2C-D**) significantly altered the avoidance phenotype, nor did it significantly affect the proportion of choosers committing to a particular OP50 lawn (**Figure S2E-F**). Only L1 larvae failed to exhibit corpse lysate-induced aversion (**Figure 2C**) whereas choosers at L2 and older were repelled by the presence of worm corpse lysates (**Figure S2G**). Lysates of inactivated worm eggs failed to promote an aversive response (**Figure S2H**), but lysates of L1 worms and older triggered aversion (**Figure S2I**). Using starved worms as well as replacing OP50 with *Comamonas* DA1877 as an alternative food source, we further confirmed that the aversive behavior is independent of feeding status and bacterial food source (**Figure 2D-F**). Adult males were equally responsive to the aversive cue elicited by worm corpse lysates as hermaphroditic choosers (**Figure 2G**). Taken as a whole, these results suggest that the avoidance of dead conspecifics represents a fundamental behavioral response in *C. elegans*, that is not confounded by age, nutritional status, or sex.

**Figure 2.**
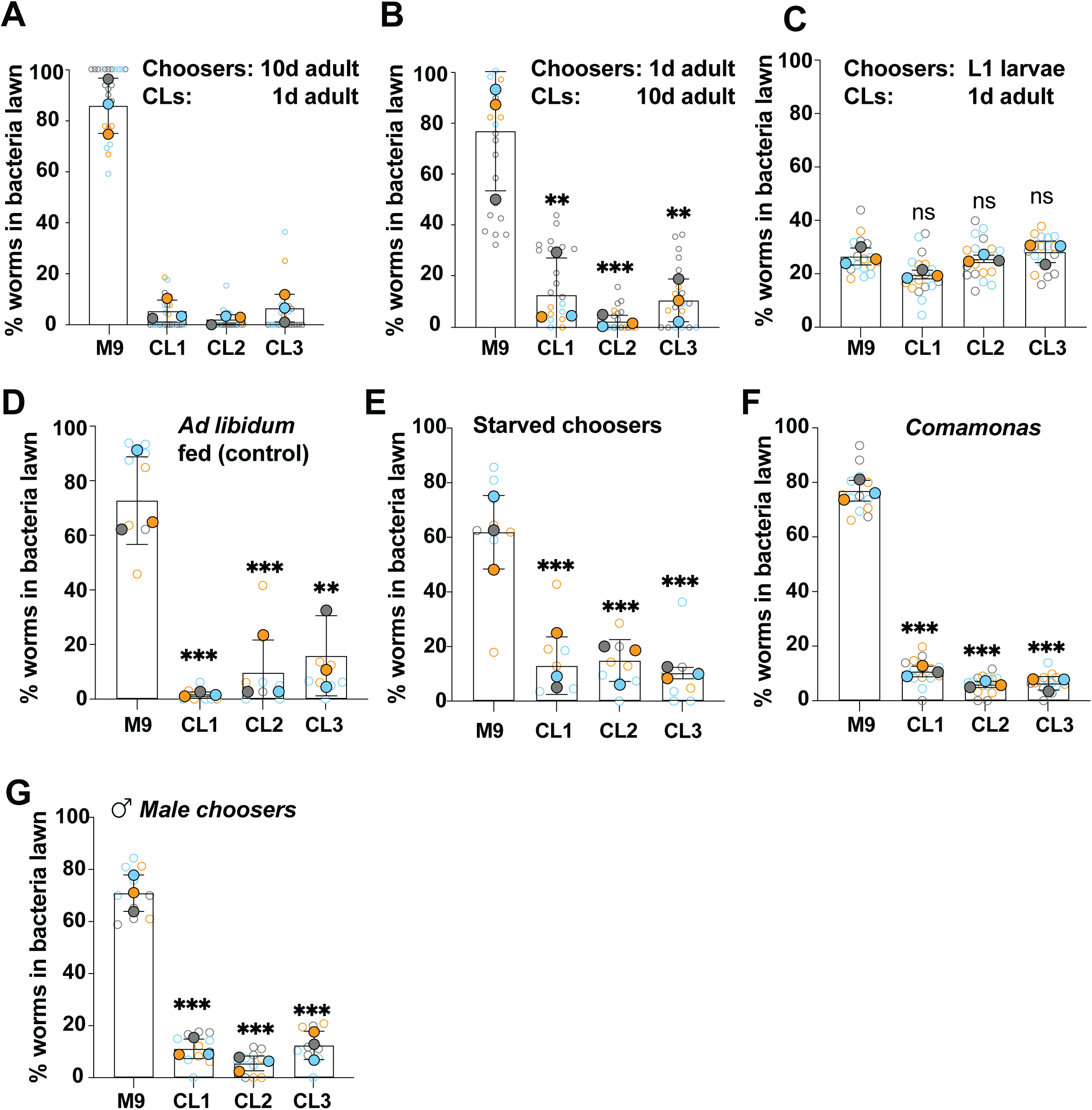
Factors influencing the aversive behavior. (A-G) Four-option choice assays testing worm behavior upon exposure to worm corpse lysates. In each experiment, three *E.coli* OP50 lawns were supplemented with 10μg worm lysate whereas the fourth *E.coli* OP50 lawn was supplemented with M9 and served as control. (A) Assays testing 10-day old choosers and corpse lysates of 1-day old adults. (B) Assays testing 1-day old choosers and corpse lysates of 10-day old adults. (C) Assays testing L1 larval choosers and corpse lysates of 1-day old adults. (D-E) Assays testing 1-day old fed (D) or starved (E) choosers and corpse lysates of 1-day old adults. (F) Standard choice assay in which *E.coli* OP50 lawns were replaced with *Comamonas* spp. lawns. (G) Standard choice assay performed with male choosers. For (A) through (G): CL1: lysate of NaN_3_-inactivated worms; CL2: lysate of worms lysed without prior inactivation; CL3: lysate of ethanol-inactivated worms. Solid circles represent individual biological replicates; hollow circles represent individual technical replicates; corresponding biological and technical replicates are color matched. Error bars represent standard deviation of the mean. Indicated P values were calculated using 1-way ANOVA tests with multiple comparison, with M9 control condition serving as reference. ns=p>0.05 (not significant); *p=<0.05; **p=<0.01; ***p=<0.001.

### Death perception impacts fitness and reproduction

Previous work in the vinegar fly *Drosophila melanogaster* demonstrated that the exposure to dead conspecifics significantly shortened fly lifespan^5,6^. We thus sought to determine whether death perception alters physiology and longevity in exposed worms. Assessing worm survival in lifespan assays, we found that the exposure to worm corpses significantly reduced *C. elegans* lifespan (**Figure 3A**). In thrashing assays, which assess nematode motility and health, we further observed that adult worms previously exposed to worm corpse lysates showed a significant decrease in thrashing rate, indicative of reduced fitness (**Figure 3B**). The exposure to worm corpse lysates also led to a short-term increase in egg laying in young adults during the first 24 hours of exposure (**Figure 3C**). These results show that the perception of dead conspecifics has a profound impact on worm physiology and reproduction.

**Figure 3.**
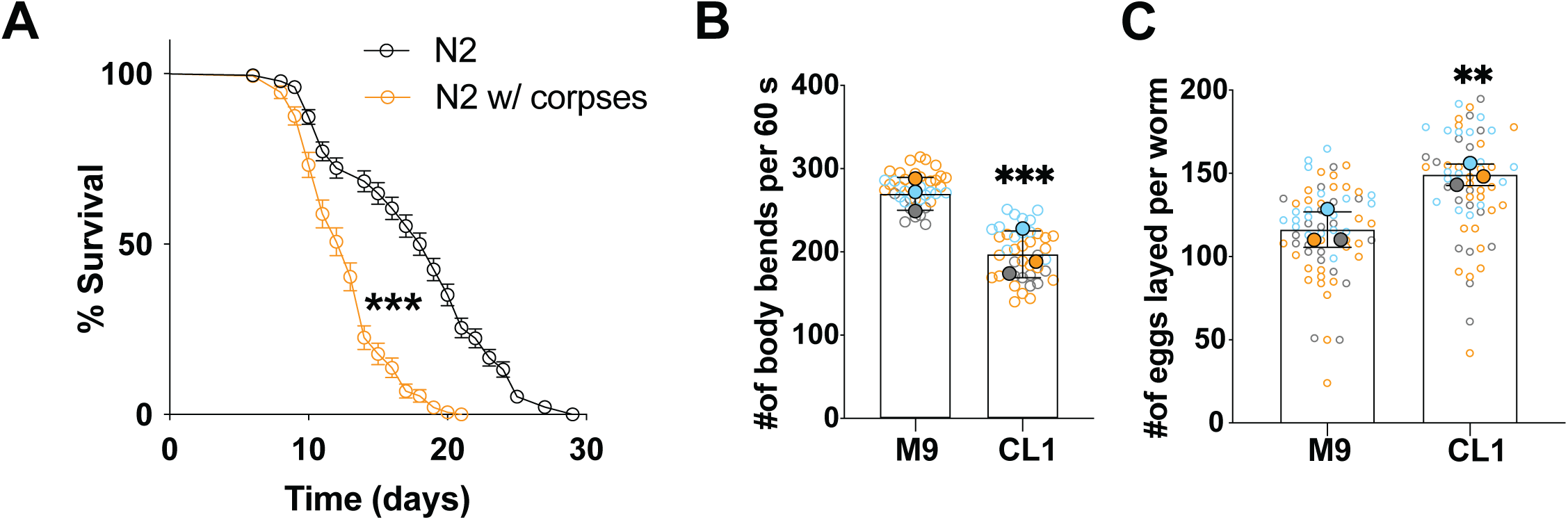
Death perception impacts fitness and reproduction. (A) Lifespan of *C. elegans* in the presence (orange) or absence (black) of naturally deceased worms (n=228 (control) and 146 (w/ corpses)). (B) Thrashing assay to determine worm fitness. CL1: lysate of NaN_3_-inactivated worms. (C) Quantification of egg laying in the presence or absence of worm corpse lysates. For (B-C): Solid circles represent individual biological replicates; hollow circle represent individually assessed animals. Corresponding biological and technical replicates are color matched. Error bars represent standard deviation of the mean. Indicated P values were calculated using Mantel-Cox test (A) and unpaired two-sided t tests (B-C). ns=p>0.05 (not significant); *p=<0.05; **p=<0.01; ***p=<0.001.

### Olfactory neurons AWB and ASH are required for death perception-induced behavioral and physiological changes

Our initial experiments showed that similar quantities of intact corpses and corpse lysates were equally potent to promote aversive behavior in choosers (**Figure 1**). We thus hypothesized that death perception was based on chemosensory cues, rather than visual or mechanosensory inputs. Indeed, we found that *che-2* and *che-3* deficient animals, which have severe defects in chemosensory ciliated neurons^20,21^, did not avoid corpse lysate spotted bacterial lawns (**Figure 4A, S4B**). Next, we performed four-option choice assays in which worm corpse lysates were applied to cotton swabs fixed to assay plate lids, hovering over the individual bacterial lawns without touching them (**Figures 4B**). We found that the presence of a corpse lysate-soaked cotton swab was sufficient to provoke aversive behavior in choosers (**Figure 4C**), suggesting that an olfactory, rather than a gustatory cue is required to detect dead conspecifics. We confirmed these results using an alternative experimental setup in which corpse lysates were applied on agar pedestals hovering over the bacterial lawns (**Figure S3A-B**). Based on these results, we predicted that death perception is mediated by chemosensory neurons responsive to volatile cues. In *C. elegans*, the main chemosensory organ consists of twelve amphid and one phasmid neuron pair that perceive external and internal signals^22–25^. The dendrites of these sensory neurons terminate in cilia that are directly exposed to the environment^26^. To identify the neurons required for death perception, we tested single neuron ablation strains of amphid neurons critical for chemoattraction (ASE, ASG, ASI, ASK, AWA, AWC, BAG), chemorepulsion (ASH, ADL, AWB), and dauer formation (ADF, ASJ)^17^ in choice assays. Our results showed that AWB and ASH neurons are required for death perception-mediated aversion in all tested four-option choice assays paradigms (**Figure 4D-E, S3B-G**). In contrast, all other evaluated amphid neurons were dispensable for the death perception process (**Figure S4A-M**). Lifespan and thrashing assays further confirmed that AWB and ASH neurons are necessary to induce physiological changes in response to death perception (**Figures 4F-I, S3H-J**). These results demonstrate that the perception of dead conspecifics in *C. elegans* is mediated by the olfactory perception of a volatile cue through AWB and ASH neurons.

**Figure 4.**
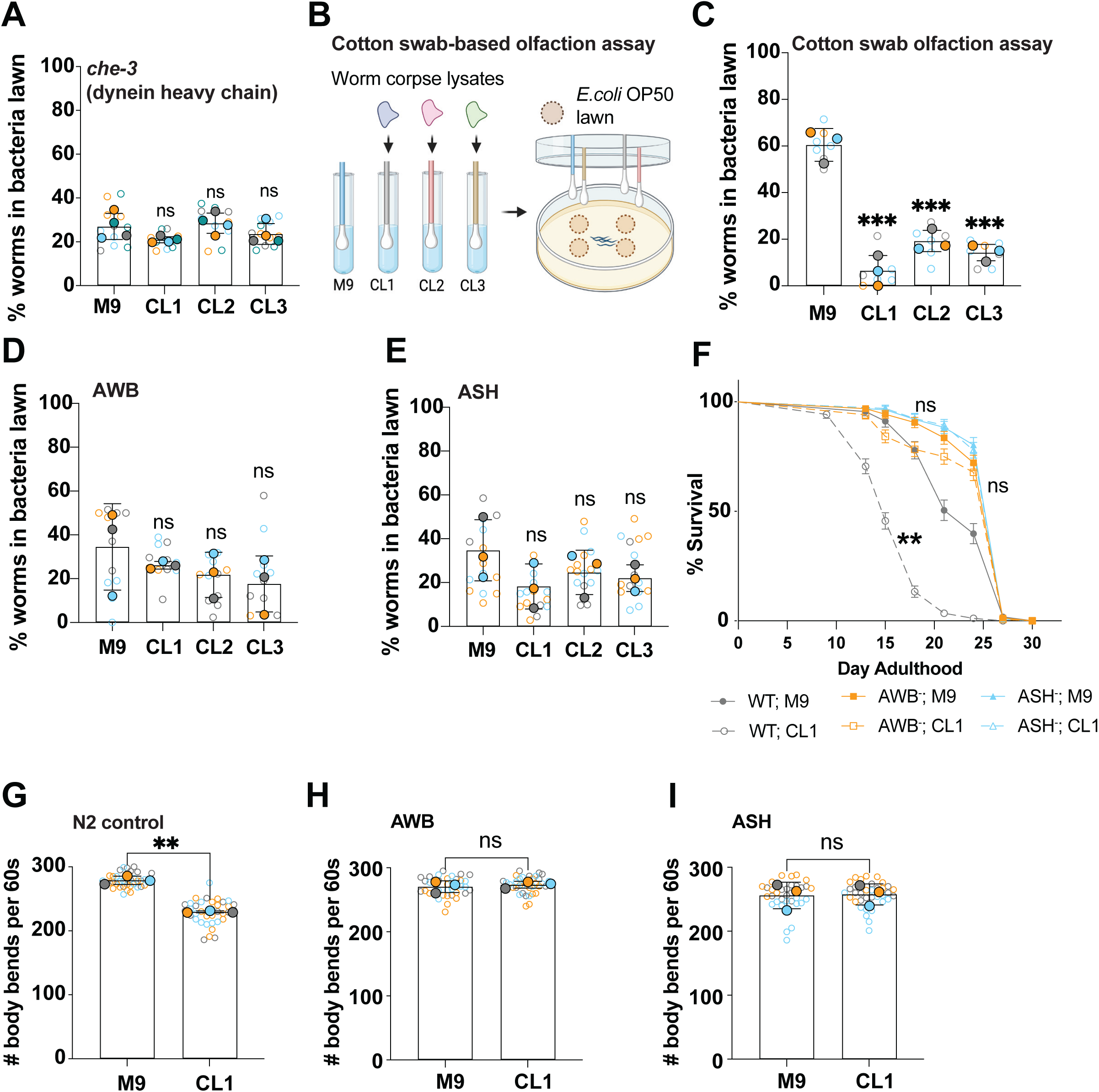
Olfactory neurons AWB and ASH are required for dead perception-induced behavioral and physiological changes. (A) Four-option choice assays testing behavior of *che-3*(*e1124*) animals upon exposure to worm corpse lysates. In each experiment, three *E.coli* OP50 lawns were supplemented with 10μg worm lysate whereas the fourth *E.coli* OP50 lawn was supplemented with M9 and served as control. (B) cotton swabs-based assay schematic. (C) Four-option choice assays in which choosers were exposed to three cotton swabs soaked with 20μl worm corpse lysate (approximate protein concentration in lysates: 1mg/ml) and one cotton swap containing an equal volume of M9. Q-tips were hovering over *E.coli* OP50 lawns without contacting them. (D-E) Four-option choice assays testing worm behavior of AWB-(D) or ASH-deficient (E) worms upon exposure to indicated worm corpse lysates. CL1: lysate of NaN_3_-inactivated worms; CL2: lysate of worms lysed without prior inactivation; CL3: lysate of ethanol-inactivated worms. (F) Lifespan of indicated *C. elegans* strains in the presence (CL) or absence (M9) of worm lysates. (G-I) Thrashing assays of 1-day adult N2 wildtype (G) AWB-deficient (H) and ASH-deficient (I) worms in the presence (CL) or absence (M9) of worm corpse lysates. For (A, C-E, G-I): Solid circles represent individual biological replicates; hollow circles represent individual technical replicates (C-E) or individually assessed animals (G-I). Corresponding biological and technical replicates are color matched. Error bars represent standard deviation of the mean. Indicated P values were calculated using 1-way ANOVA tests with multiple comparison with M9 control condition serving as reference. (A-E), Mantel-Cox regression analysis (F), and unpaired two-sided t tests (G-I). ns=p>0.05 (not significant); *p=<0.05; **p=<0.01; ***p=<0.001.

### Death perception engages glutaminergic signaling and involves the guanylate cyclase, DAF-11, and the TAX-2/TAX-4 cGAMP-coupled GPCRs

When responding to environmental cues, *C. elegans* olfactory neurons engage complex signaling cascades to relay information to deeper neuronal layers (**Figure 5A-C**). These neurons serve to process and integrate external information and initiate the execution of a motor program^27^. To start understanding the neuronal circuitry and signaling modalities involved in death perception, we tested a panel of *C. elegans* mutant strains deficient in specific interneurons, neurotransmitters, G-protein coupled receptors (GPCRs), voltage-gated channels, and transient receptor potential (TRP) cation channels, in four-option choice assays (**Figure 5D-H**, **Figure S5A-U**). We found that *unc-25* and *unc-47* worms deficient in GABAergic signaling (**Figure S5B-C**) as well as worms incapable of synthesizing octopamine, dopamine, or serotonin (**Figure S5D-J**) continued to avoid bacterial lawns spotted with worm corpse lysates, suggesting that GABAergic, dopaminergic, serotonergic, and octopamine-mediated signaling is dispensable for death perception. Contrasting, worms with a loss-of-function mutation in the glutamate transporter, *eat-4*, failed to avoid worm corpse lysates (**Figure 5D, S5K**). We further identified the guanylate cyclase, *daf-11,* the *tax-2*/*tax-4*-containing GPCR (both expressed in AWB neurons), and the cGMP-dependent kinase, *egl-4* (expressed in ASH neurons) as necessary for death perception (**Figure 5E-H**). The ablation of *ocr-2* and *osm-9* TRPV channel subunits required in ASH neurons for the promotion of behavioral changes in response to food shortage and increased population density^28,29^ did not prevent an aversive response to the presence of worm corpse lysates (**Figure S5L-M**). Death perception thus represents a rare example in which ASH activation does not involve OCR-2/OSM-9 TPRV channels^30^. In additional experiments, we found that deficiencies in neuroligin (*nlg-1*), which inhibit a subset of sensory behaviors^31–34^ did not prevent corpse lysate avoidance (**Figure S5N**), neither did the expression of loss-of-function alleles of the *odr-1*-type guanylate cyclase, which is essential for AWC-mediated olfaction and odor discrimination^35,36^ (**Figure S5O**). Notably, the ablation of AIA, AIB, and AIY interneurons individually or combinatorially did not interfere with corpse lysate-associated aversion, (**Figure P-T**), suggesting the involvement of a neuronal circuit relying on AIZ or independent of the first interneuron layer.

**Figure 5.**
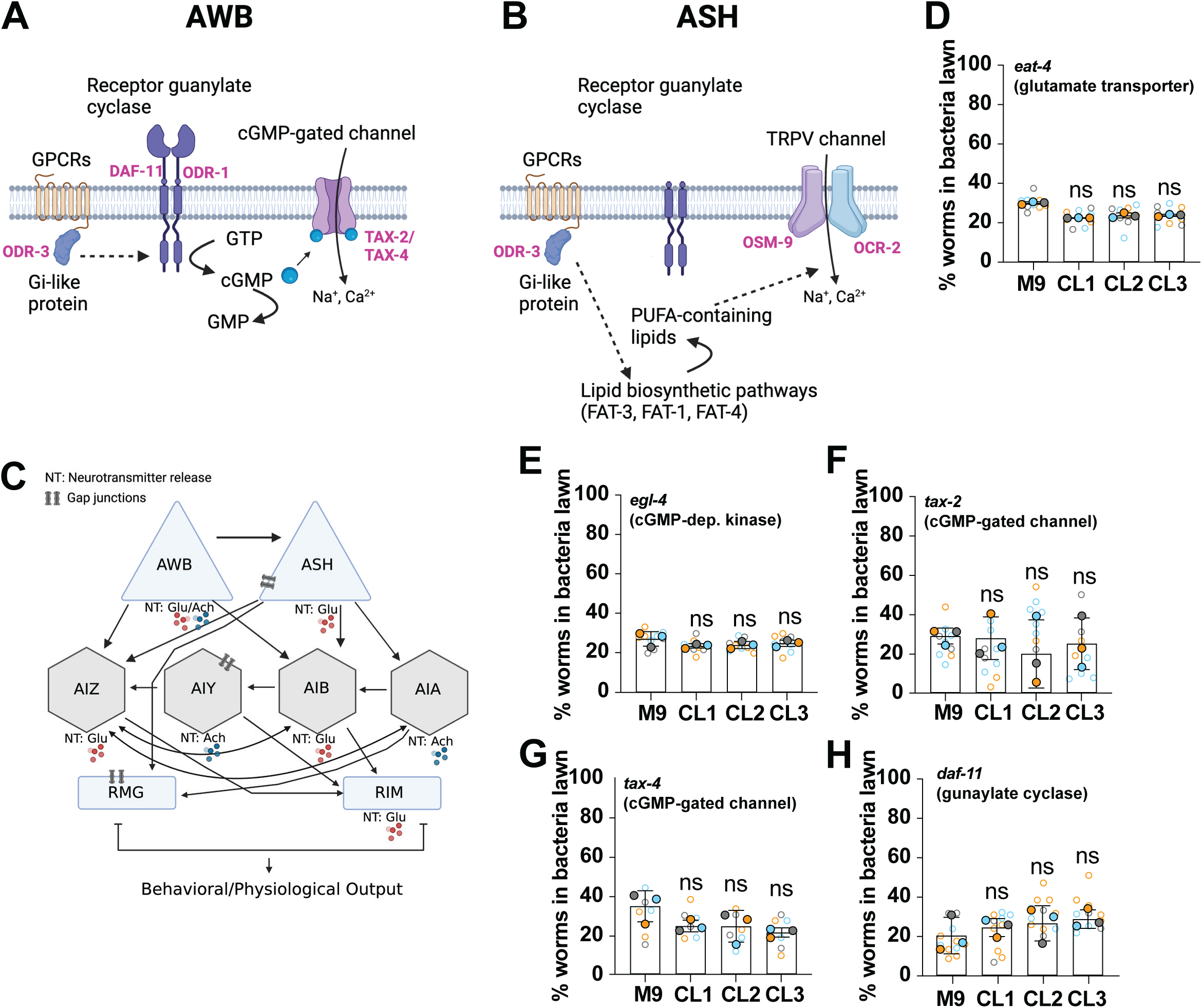
Death perception involves glutaminergic signaling, GPCR activity, and cGMP-gated channels. (A-B) Schematics of key signaling proteins involved in AWB (A) or ASH (B) function. (C) Circuit diagram of how AWB and ASH are proposed to signal to inter- and motor neurons. (D-H) Four-option choice assays testing worm behavior of strains deficient in *eat-4* (D), *egl-4* (E), *tax-2* (F), *tax-4* (G), or *daf-11* (H) worms upon exposure to indicated worm corpse lysates. CL1: lysate of NaN_3_-inactivated worms; CL2: lysate of worms lysed without prior inactivation; CL3: lysate of ethanol-inactivated worms. Solid circles represent individual biological replicates; hollow circles represent individual technical replicates. Corresponding biological and technical replicates are color matched. Error bars represent standard deviation of the mean. Indicated P values were calculated using 1-way ANOVA tests with multiple comparison with M9 control condition serving as reference. ns=p>0.05 (not significant); *p=<0.05; **p=<0.01; ***p=<0.001.

### AMP and Cysteine serve as potential death cues recognized by *C. elegans*

Death perception is a sensory modality observed in most metazoan animals. We thus hypothesized that the presence of a conserved, yet rudimentary death signature could be driving this response. To test this prediction, we exposed *C. elegans* to lysates of dead *C. briggse* and *C. remani* – two close relatives of *C. elegans* – as well as dead vinegar flies or planarians (**Figure 6A-D, S6A-B**). We found that all corpse lysates were effective at inducing aversion in four-option choice assays, supporting our hypothesis. We also excluded the involvement of a secreted factor from *C. elegans* in this process as conditioned M9 buffer failed to introduce an aversive response in the utilized choice paradigm (**Figure S6C**). We then sought to further understand the biophysical and biochemical nature of the death cues. We observed that the experimental procedure used to kill worms prior to lysis directly impacted how effective worm corpse lysates were at inducing an aversive response. Lysates of heat-killed (65°C, 1hour) worms (CL5) did not trigger avoidance behavior (**Figure S6D**). Contrasting, lysates of worms homogenized without prior killing (CL2) or treated with 100% ethanol prior to lysis (CL3) promoted aversion (**Figure 6A**). Aversive responses tended to be strongest for worm lysates derived from animals killed with the mitochondrial complex IV inhibitors, sodium azide (NaN_3_; CL1) or potassium cyanide (KCN; CL4) (**Figure 6E**). Neither repeated freeze-thawing (**Figure S6E**), boiling of corpse lysates for 1 hour (**Figure S6F**), incubating lysates for several weeks at room temperature (**Figure S6G**), nor treatment of lysates with DNAase (**Figure S6H**) or proteinase K (**Figure S6I**) diminished the lysate’s potency to promote aversion. This suggests that the death cues are relatively stable metabolites responsive in concentration to mitochondrial activity and depleted in worms dying of heat stress. We, therefore, used polarity-based fractionation to separate *C. elegans* corpse lysates into less-complex fractions. In choice assays, we identified the extract fraction which induced the strongest avoidance response (**Figure S6J**) and analyzed this fraction, as well as unfractionated corpse lysates, using mass spectrometry (MS) and nuclear magnetic resonance (NMR). These experiments identified a total of 52 unique metabolites across samples (**Supp. Tables S1-S3**). The NMR-based analysis identified several amino acids, including alanine, histidine, cysteine, and glycine, and the nucleotide adenosine mono-phosphate (AMP) as shared between the tested samples. Contrasting, MS identified several phosphocholines, phosphoethanolamines, amino acids, and nucleotides, including AMP, as constituents of the samples, only four of which were present in both the active complete lysate (CL1) and the active fractionated lysate (**Figure S6K-L**, **Table S3**). We then tested several of these compounds individually in four-option choice assays to determine their ability to induce aversive behaviors. Of particular interest was the nucleotide AMP, which was the only metabolite identified in both complete and fractionated lysates analyzed by NMR or MS as well as the amino acids alanine, histidine, arginine, and cysteine, which were abundantly present in the active worm corpse lysate fractions. Using our four-option choice paradigm, we found that adding 100 μM AMP or cysteine to inactive CL5 corpse lysate was sufficient to render it into a potent repellant (**Figure 6F-G**). In contrast, supplementing CL5 with up to 100 mM glucose or lactate, two metabolites we identified as constituents of the active corpse lysate fraction (**Supp. Tables S1-S2**), did not convert CL5 into an aversion-promoting signal (**Figure S7A-B**). Ectopic supplementation of a single bacterial food lawn with 10 μl of 100mM cysteine and histidine, but not alanine, glycine, arginine, glucose, or AMP was sufficient to promote aversion in adult worms (**Figure S7C-F**). Taken as a whole, our results are consistent with a model in which the intracellular metabolites, AMP and cysteine are recognized by *C. elegans,* possibly in concert with additional molecules, as a death signature when present in the environment.

**Figure 6.**
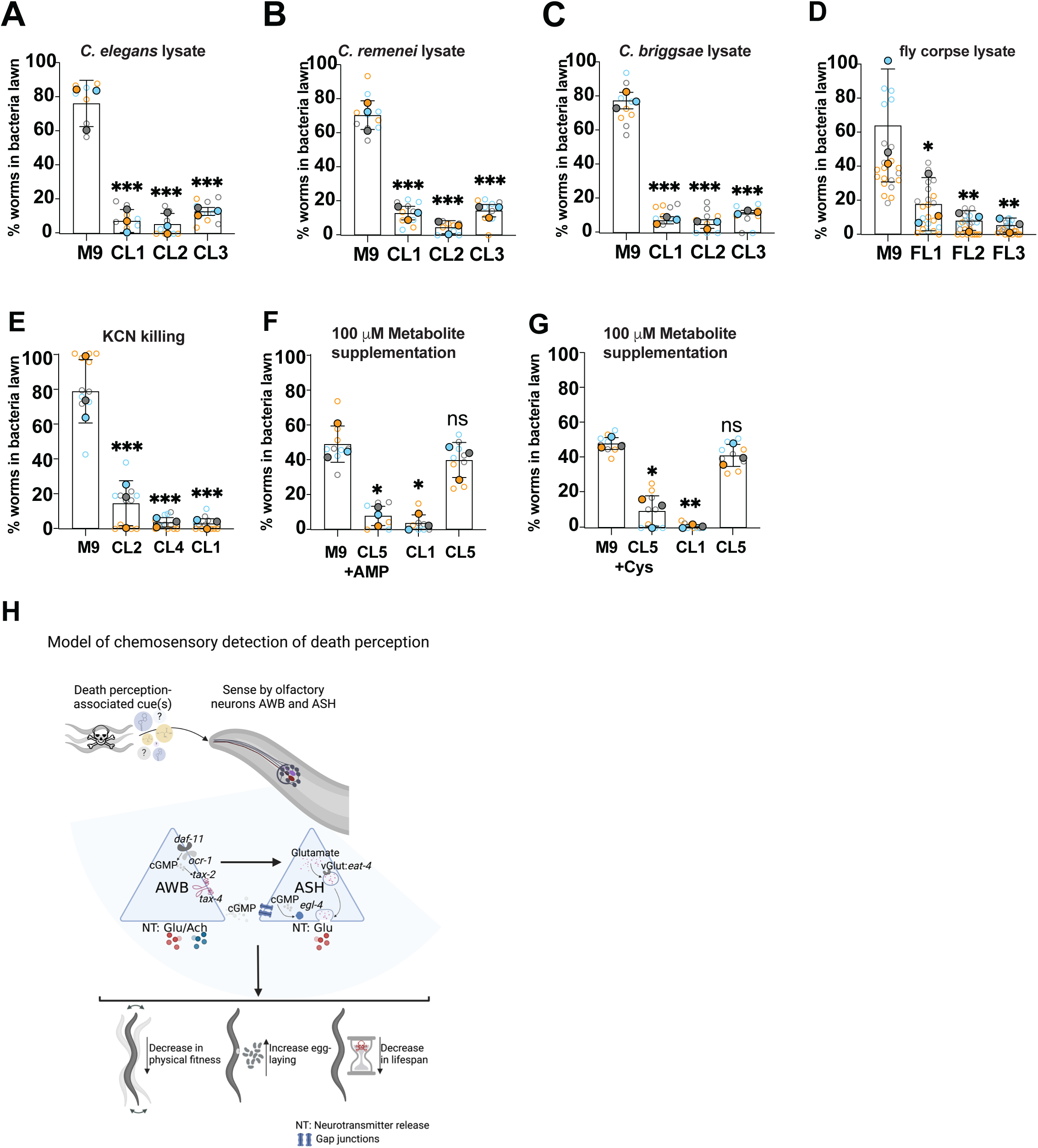
The intracellular metabolites cysteine and ATP are abundant in corpse lysates and sufficient to trigger avoidance behavior. (A-G) Four-option choice assays in which 1-day old adult choosers were exposed to indicated corpse lysates of chemically inactivated *C. elegans* (A), *C. remenei* (B) *C briggsae* (isolate 1977) (C), and *Drosophila melanogaster* (vinegar fly) (D). For (E), worms were inactivated with either NaN3 (CL1), KCN (CL4 KCN), or homogenized without prior inactivation (CL2). For (F-G), *E.coli* OP50 lawns were supplemented either with lysates of NaN_3_ inactivated worms (CL1), M9 buffer, lysate of heat-killed worms (CL5), or CL5 supplemented with 100 μM AMP (F) or cysteine (Cys) (G). (H) Schematic representation of key findings of this study. NT: Neurotransmitter; Glu: Glutamate; Ach; Acetylcholine For (A-G): Solid circles represent individual biological replicates; hollow circles represent individual technical replicates. Corresponding biological and technical replicates are color matched. Error bars represent standard deviation of the mean. Indicated P values were calculated using 1-way ANOVA tests with multiple comparison with M9 control condition serving as reference. ns=p>0.05 (not significant); *p=<0.05; **p=<0.01; ***p=<0.001.

## Discussion

All organisms respond to a rich compendium of both simple (e.g. smell, taste, mechanosensation) and more complex (crowding, hunger/thirst, sexual attraction) sensory inputs, shaping their immediate and long-term behavior. Many of these cues have a species-specific impact on health, vitality, and lifespan. The recognition of dead conspecifics and corpses of unrelated species is an ancient skill possessed by most metazoans. The presence of non-related cadavers and deceased prey results in species-specific behaviors and neuronal states which may include fear, avoidance, exploratory curiosity, and hunger/urge to eat^13^. The recognition of dead conspecifics, however, frequently signals the presence of imminent danger and/or the lack of essential resources in the immediate environment. Reflective of the evolutionary history and occupied ecological niche, the response to the presence of dead conspecifics may consist of a species-specific behavioral change. Our work identified the nucleotide AMP, and the amino acid cysteine, as potential cues involved in death perception in *C. elegans*. We also show that that corpse lysates of unrelated species trigger avoidance behavior in *C. elegans*. This may well reflect a species-specific cue-response paradigm. However, it is interesting to consider that, as they disintegrate, apoptotic human cells release a “metabolite secretome”, which is recognized by neighboring tissue as a death signature, resulting in transcriptional changes^37^. This secretome consists of simple molecules associated with the intracellular space, including GMP, IMP, spermidine, and UDP-glucose. We thus speculate that, while the cues are likely species-specific, the concept of detecting a death signature consisting of a mixture of intracellular metabolites might well be conserved. Our results support a model (**Figure 6E**) in which such a death signature present in the environment is sufficient to trigger a behavioral response in *C. elegans*, mediated by two pairs of sensory neurons that directly sample the environment. Whether or not this mechanism extends beyond the recognition of apoptotic neighboring cells and tissues in mammals remains to be tested.

The influence of sensory perception on aging and health is well described in *C. elegans*. Mutants with defects in sensory cilia or sensory signal transduction are long-lived^38^. The ablation of specific olfactory (AWA, AWC) or gustatory neurons (ASI, ASG) increases lifespan by engaging the insulin/IGF-1-like (*daf-2*/*daf-16*) and SKN-1 signaling pathways^17,39^. Recent studies have further shown that socio-environmental cues, such as crowding or the presence of injured conspecifics, affect *C. elegans* physiology and fitness^40,41,42^. The perception of injured conspecifics is mediated by amphid sensory neurons ASI and ASK and modulated by neurotransmitters GABA and serotonin. While promoting aversive behavior, the recognition of injured conspecifics does not affect lifespan. Our results show that AWB and ASH-mediated death perception in *C. elegans* shortens lifespan, induces aversive behavior, and affects reproductive fitness. The finding that the exposure to dead conspecifics leads to an acute increase in egg laying may result as a secondary outcome of amphid neuron activation. Indeed, exposure to the noxious stimulus, Cu^2+^ leads to an ASH-dependent short-term increase in egg laying rates before collapsing back to or below control levels^43,44^, demonstrating that ASH activation directly affects reproductive aging.

Perhaps the most important finding of our work is the identification of a novel cue-neuron-behavior paradigm that allows us in future experiments to interrogate and manipulate individual aspects of this system to better understand how sensory inputs shape physiological and behavioral outputs. We acknowledge that species-specific evolutionary needs and adaptations to ecological niches make it less likely for such cue-neuron-behavior paradigm to be conserved and directly translate into more complex organisms, including humans. Nevertheless, this study provides novel information on how input through sensory neurons shapes internal states, a concept that is also applicable to human behavior and health.

Our work leaves us with several interesting questions: What are the molecular mechanisms that translate death perception into physiological changes? Could simple metabolic signatures serve as conserved death cues across species? And what is the neuronal circuitry orchestrating the signal transfer from sensory neurons to the periphery? In answering these questions in future studies, we will obtain a comprehensive picture of how this modality of death perception is mediated in metazoans.

### Limitation of this study

The parallel loss of multiple sensory neurons is lethal in *C. elegans*, which prevents us from testing whether AWB and ASH neurons are sufficient for death perception. Technical limitations to deplete AMP and cysteine in worm corpse lysates further prevents us from testing if either molecule is required for corpse lysates to trigger avoidance.

## Author contributions

MCT supervised the project. MCT and MHL planned and designed the experiments. MHL, BS, NDU, and MCT performed all experiments. MCT and MHL wrote the manuscript. All authors edited and approved the final manuscript. All roles and responsibilities within the project team were agreed upon amongst contributors ahead of the research.

## Data availability statement

Unprocessed numeric datasets (excel tables and prism files) containing and analyzing the data presented in this study are available from the corresponding author upon reasonable request.

## Acknowledgments

We thank the members of the Truttmann lab for helpful for proof-reading manuscript drafts, comments and discussion. Ashootosh Tripathi and Fei Yang from the Natural Products Discovery Core at University of Michigan are acknowledged for help with sample fractionation and cue identification. David Paris assisted in the preparation of worm plates used for olfactory tests. Kristi Gendron provided vinegar fly lysates and Longhua Guo planaria lysates. We also thank the *Caenorhabditis* Genetics Center, which is funded by NIH Office of Research Infrastructure Programs (P40 OD010440), and Drs. Denise Ferkey, Cori Bergmann, Dennis Kim, and Yuichi Lino for sharing strains. MHL is a Rackham fellow and obtained support from NIA Training Grant AG000114, 1F31DC02039701 (Kirschstein NRSA), and 1F99NS135768 (D-SPAN). NDU was supported by training grants GM008322, AG000114 and a Kirschstein NRSA (1F31AG08589101). MCT is supported by grant 1R35GM142561. MHL and MCT are responsible for the data presented in this publication. The presented findings are not necessarily representative of the official views of the National Institutes of Health.

## Material and Methods

### Strain maintenance

All *Caenorhabditis* strains used in this study were cultured on nematode growth media (NGM) plates at 20°C using *E. coli* OP50 bacteria as food source and following standard procedures^45^. The Bristol N2 strain served as wildtype (WT) reference in all experiments. Additional strains used in this study were obtained from the *Caenorhabditis* Genetics Center (University of Minnesota, http://www.cbs.umn.edu/CGC/) or the scientific community and outcrossed into N2 Bristol as needed^46–48^. All experiments were conducted using synchronized worm populations obtained by hypochlorite treatment (hypochlorite bleaching buffer: [56.6ml of distilled water with 14.4mL of 5N NaOH and 6.6mL of 8.25%NaHOCl]) unless otherwise stated. **Supplementary Table S4** lists all *Caenorhabditis* strains used in this study.

### Culturing of bacteria

*E. coli* OP50 was cultured overnight in Luria Broth supplemented with streptomycin (LB media) shaking (250 rpm) at 37 °C. HT115 was cultured under the same conditions, except LB was supplemented with carbenicillin, while Comamonas (DA1877) and *Pseudomonas aeruginosa* PA14 (PA14) were cultured overnight in LB media without antibiotics.

### General assay considerations

All assays were performed with synchronized, day 1 adult hermaphroditic *C. elegans* animals at 20°C unless otherwise stated.

### PA14 Avoidance assays

7 µL of an overnight LB culture of PA14 was spotted onto 35mm SKA plates (SKA plates: 3 g/L NaCl, 3.5 g/L peptone, 17 g/L agar, 5mg/L cholesterol in ethanol, 1 mL/L 1 M CaCl_2_, 1 mL/L 1 M MgSO_4_, and 25 mL/L 1 M potassium phosphate buffer (pH 6.0)). The plates were then incubated at 37°C for 24 hours to allow the PA14 lawn to grow, after which they were transferred to 20°C for an additional 24 hours before usage. Assays were scored after day 1 adult worms were allowed to choose for 3 hours at 25°C.

### Corpse and corpse lysate avoidance assays

35 mm NGM plates were inoculated with 50 µl of *E. coli* OP50 to grow a central, single bacterial lawn. Next, the lawn was supplemented with approximately 60 intact *C.elegans* corpses or 10 µg corpse lysate in M9. Approximately 30 animals (choosers) were then placed in the center of the plate. After 3 hours, the number of animals remaining in the lawn and occupying the periphery were counted.

### Four-option Choice Assays

35mm NGM plates were divided into 4 equal quadrants. Each quadrant was inoculated with 10 µl *E. coli* OP50 to grow four equally distributed bacterial lawns. In each assay, approximately 30 animals (choosers) were placed in the center of the plate. Lawn occupation was quantified after three hours. Unless specified otherwise, four-option choice assays were performed using hermaphroditic day 1 adult animals as choosers and corpse lysates of chemically inactivated day 1 adults as repellants. In standard experiments, three of the four *E.coli* OP50 lawns were supplemented with approximately 10μg worm corpse lysate in 10µL of M9 and a one lawn with an equal volume of M9 prior to assay start. Lawn occupation was quantified after 3 hours. Animals not committed to any bacterial lawn after 3 hours were designated as non-choosers. Experiments involving *P. aeruginosa* (P.14) used 35mm SKA plates spotted with 3 µL of *Pseudomonas aeruginosa* isolate PA14 (PA14) in one quadrant, and 7 µL of *E. coli* OP50 in the other three quadrants. Assays testing starved choosers were performed using M9-washed day 1 adult animals previously kept on bacteria-free 100 mm NGM plates for 24h.

### Preparation of Worm, fly, and planaria lysates

Synchronized worm populations of specified ages were collected and washed three times with M9. Animals were then treated with 1000 µL 1M Sodium Azide (NaN_3_), 1000 µL M9, or 1000 µL 100% Ethanol for 1 hour. After this inactivation step, animals were washed five times with 1000 µL M9 and pelleted by centrifugation for 30 seconds at 3,000 g after each wash. Samples were then and homogenized using a Qiagen Tissue lyser II for 10 min at 30 Hz at 4°C. The corpse homogenate was centrifuged for 2 min at 13,000g and the supernatant subsequently filtered using 0.22 µM filter. Protein concentrations of cleared lysates were determined using a BCA assay (Pierce). Lysates were adjusted with M9 to a protein content of 1 µg/µl and tested in the 4-option choice assay. In all figures, the different lysate preparations are referred to as follows: corpse lysate 1 (CL1): lysate of NaN_3_-inactivated worms; corpse lysate 2 (CL2): lysate of worms lysed without prior inactivation; corpse lysate 3 (CL3): lysate of Ethanol-inactivated worms. Protein concentrations of cleared lysates was determined using a microBCA assay (Pierce). To test the potential involvement of a molecule secreted by *C. elegans* in avoidance behavior, conditioned M9 buffer, in which synchronized day 1 adult worms were kept for 1 hour, was used.

For fly lysate preparations, Canton S embryos were collected using PBS and placed at equal numbers into standard cornmeal-sugar-yeast medium at 25°C with 12:12-hour light:dark cycles and 60% relative humidity. The resultant adult flies were collected within 24 hours of emergence into new bottles containing standard media and allowed to mate for 2 days. Male flies were then separated from female flies using light CO_2_ and placed into vials containing standard food that had either vehicle (100 µl of PBS) or sodium azide (1 M NaN_3_ in PBS) that was previously added to the top of the food and allowed to penetrate food overnight. After the animals were exposed to either vehicle or sodium azide for 16 hours, the flies were washed 3 times with PBS and centrifuged at 1,000g for 30 seconds. Following this, animals were homogenized in QIAGEN tissue lyser II for 10 minutes at 30Htz. The samples were then centrifuged for 1.5 minutes at 12,000g. The resulting lysate sample was filtered using a 0.22µm. Protein concentrations of cleared lysates were determined using a BCA assay (Pierce). Lysates were adjusted with M9 to a protein content of 1 µg/µl and tested in the 4-option choice assay.

For planaria lysates, *Schmidtea mediterranea* were cultured as described in Guo et al^49^. Animals were washed 3 times with PBS and centrifuged at 1,000g for 30 seconds. Following this, animals were homogenized in QIAGEN tissue lyser II for 10 minutes at 30Htz. The samples were then centrifuged for 1.5 minutes at 12,000g. The resulting lysate sample was filtered using a 0.22µm. Protein concentrations of cleared lysates were determined using a BCA assay (Pierce). Lysates were adjusted with M9 to a protein content of 1 µg/µl and tested in the 4-option choice assay.

### Long-term maintenance of large, synchronized worm populations

To collect corpses of naturally deceased worms and examine the role of age (both choosers and sample lysates) as an experimental variable, synchronized day 1 adult animals were kept on NGM plates supplemented with 1 mM IPTG and 100 μg/ml carbenicillin and nystatin, seeded with *E. coli* HT115 expressing an siRNA precursor targeting *pos-1*. POS-1 is a CCCH-type zinc finger protein almost exclusively expressed during early embryonic development. RNA interference-mediated *pos-1* knockdown prevents eggs from hatching, thereby maintaining a synchronized worm population. Corpses of naturally-deceased day 10 – day 30 worms were collected and either used immediately in four-option choice assays or stored at −80°C for future usage.

### Thrashing assays

Synchronized adult worms were cultured on 60 mm NGM plates seeded with *E. coli* OP50 until they reached day 1 of adulthood. At day 1 of adulthood, worms were washed three times with M9 and centrifuged at 1,000g for 30 seconds. Animals were then transferred to fresh *E. coli* OP50 plates treated with either 100 µL of M9 buffer or lysate from worms previously killed by 1 M sodium azide (NaN_3_) for 24 hours. After 24 hours of exposure, worms were transferred to unseeded NGM plates for 5 minutes to remove residual bacteria and then transferred to 20 µL of M9. After a 30-second acclimation period, full-body bends (thrashes) were quantified over a 60 secs. A single thrash was defined as a complete bend to one side followed by a return to the initial posture. For each biological replicate, 12–15 worms were assessed, and three biological replicates were performed per strain.

### Egg laying assay

Individual 1 day old adults were transferred onto 35mm *E.coli* OP50 plates (1 worm per plate) and exposed to either 50µL of M9 buffer (control) or lysate of animals killed with 1 M NaN_3_ added to the OP50 lawn. Number of eggs released per animal were scored after 24h at room temperature. Before quantifying the eggs laid, worms were removed and transferred to new plates. Each assay tested approximately 20-30 animals/strain/treatment and was repeated three independent times. Animals injured during transfer were excluded from analysis.

### Odor exposure assays using agar pedestals and cotton swabs

For the agar pedestal-based odor assay, small circular NMG agar pads (approximately 5 mm in diameter and 5 mm in width) were prepared. Each pad was seeded with 10 µL of *E. coli* OP50 and placed in the lid of a fresh 35 mm NMG 4-option choice plate, which was also seeded with *E. coli* OP50. Following this, the agar pads were spotted with 10 µL of either worm corpse lysates or M9 buffer as a control. Day 1 adult animals were washed and transferred to the center of the assay plate. The agar pedestals in the lid were aligned to sit above the four *E. coli* OP50 lawns on the plate. The choice responses of the Day 1 adult worms were quantified after 3 hours of exposure.

### Cotton swab olfactory-based behavioral assays

To prepare the lids of the 4-option choice assay plates, four evenly spaced holes were carefully drilled using drill bit. These holes were positioned following a template designed to align with the four *E. coli* OP50 lawns on the assay plate to ensure accurate correspondence between the lawns and the positions of the cotton swabs. Day 1 adult worms were washed and placed in the center of the assay plates to acclimate to the testing conditions prior to the addition of the cotton swab-containing lids. Each perforated lid was fitted with four cotton swab heads. The cotton swabs were then infused with 20 µL of either worm corpse lysates or M9 buffer as a control. Once the swabs were prepared, the lid was carefully placed over the 4-option assay plate, ensuring that each swab head was positioned directly above its corresponding *E. coli* OP50 lawn. This setup was crucial for maintaining consistent exposure between the worms on the plate and the odor source from the cotton swabs. Lawn occupancy was quantified 3h after initial exposure.

### Lifespan Assays

25-30 synchronized day-1 adult hermaphroditic worms were placed on 60-mm NGM RNAi plates (NGM plates supplemented with 1 mM IPTG and 100 μg/ml carbenicillin/nystatin, and seeded with *E. coli* HT115 bacteria expressing *pos-1* siRNA) to prevent egg hatching and bagging. Every three days, the plates were supplemented with either 100 µL of M9 buffer or 100 µL of filtered lysate from NaN3-inactivated worm corpses. Starting on day three, worms were transferred in three-day intervals to fresh experimental plates, timed to align with each supplementation of M9 buffer or worm lysate. Worms were scored as dead if they failed to respond to gentle taps on the head and tail with a platinum wire. Worms showing bagging or explosion through vulva phenotypes were excluded from the analysis. Lifespan studies using worm corpses followed the same protocol as described above, with the modification that approximately 100 worm corpses, killed using 1 M NaN3, were added to the experimental plates.

### Rigor and experimental statistics

To minimize experimenter bias, all experiments were performed as single-blind tests. In four-option choice assays, the experimenter was blind to the position of the distinct cues tested and the genotype of the assessed animals. In lifespan, trashing, and egg laying experiments, the experimenter was blinded to the genotypes of the tested animals. Statistical Analysis was performed using GraphPad Prism (version 10.2.2) software. Unpaired two-sided t-tests, 1-way ANOVA, and Mantel-Cox tests were performed. Figure legends specify the utilized tests for each data panel. A p value of p <0.05 was used to determine statistical significance.

### RNA interference

Knock-down of *pos-1* was induced by RNAi feeding. Knockdown efficiency was validated by qPCR (data not shown). *E.coli* growth and plate seeding was performed as previously described^50^

### Figures and graphics

Bar and dot plots were generated using GraphPad Prism (version 10.2.2). Data is represented as superblots, in which each technical replicate is represented by an empty circle and the average of each biological replicate as a color-corresponding solid circle. Schematics and graphical summaries of experimental procedures were prepared using biorender (biorender.com) and Adobe illustrator (version 26.5.3)

---

**Figure S1. *C. elegans* recognize and avoid conspecific corpses.** (A) Four-option choice assays testing behavior of day 1 adult choosers in the absence of worm corpses. All four *E.coli* OP50 lawns were supplemented with 10μl of M9 buffer. (B) Schematic representation of avoidance assay setup. (C) Avoidance assay testing behavior of day 1 adult choosers in the presence and absence of worm corpses (1:1 ratio of choosers to corpses). (D) Avoidance assay testing behavior of day 1 adult choosers in the presence of indicated ratios of choosers to worm corpses. (E) Avoidance assay testing behavior of day 1 adult choosers in the presence of worm corpse lysates (CL1, CL2) and *Pseudomonas aeruginosa* isolate PA14. For (A) through (E): Solid circles represent individual biological replicates; hollow circles represent individual technical replicates; corresponding biological and technical replicates are color matched. Error bars represent standard deviation of the mean. Indicated P values were calculated using unpaired two-sided t tests (B) and 1-way ANOVA tests with multiple comparison (A, D, and E with M9 control condition serving as reference). ns=p>0.05 (not significant); *p=<0.05; **p=<0.01; ***p=<0.001.

**Figure S2. Factors influencing the aversive behavior.** (A-D, G, H) Four-option choice assays testing worm behavior upon exposure to worm corpse lysates. In each experiment, three *E.coli* OP50 lawns were supplemented with 10μl worm lysate (approximate protein concentration: 1mg/ml) whereas the fourth *E.coli* OP50 lawn was supplemented with 10μl M9 and served as control. (A-B) Assays testing 1-day old (A) and 5-day old (B) choosers reacting to corpse lysates of 1-day old adults. (C-D) Assays testing 1-day old choosers reacting to corpse lysates of 1-day old (C) and 5-day old (D) adults. (E-F) Percentage of choosers committing to a bacterial lawn by the end of the experiment. Indicated ages are days of adulthood (G) Assays testing L2 larval choosers and corpse lysates of 1-day old adults. (H) Choice assay testing behavior of day 1 old choosers in the presence of three *E.coli* OP50 lawns spiked with 10μl egg lysate (approximate protein concentration: 1mg/ml). The fourth lawn was spiked with 10μl M9 buffer. For (A) through (H): Solid circles represent individual biological replicates; hollow circles represent individual technical replicates; corresponding biological and technical replicates are color matched. Error bars represent standard deviation of the mean. Indicated P values were calculated using 1-way ANOVA tests with multiple comparison, with M9 control condition serving as reference. ns=p>0.05 (not significant); *p=<0.05; **p=<0.01; ***p=<0.001.

**Figure S3. Olfactory neurons AWB and ASH are required for dead perception-induced behavioral and physiological changes.** (A) Assay schematic of agar pedestal-based assay. Agar pedestals hover over *E.coli* OP50 lawns without directly contacting them. (B-D) Four-option choice assays in which N2 wildtype (B), AWB-deficient (C) and ASH-deficient (D) day 1 adult choosers were exposed to three agar pedestals supplemented with 10μl day 1 adult corpse lysate (approximate protein concentration: 1mg/ml) and one pedestal supplemented with an equal volume of M9. Pedestals were hovering over *E.coli* OP50 lawns without contacting them. (E-G). Four-option choice assays in which N2 wildtype (E), AWB-deficient (F) and ASH-deficient (G) day 1 adult choosers were exposed to three cotton swabs soaked with 10μl day 1 adult corpse lysate (approximate protein concentration: 1mg/ml) and one cotton swab soaked with an equal volume of M9. Cotton swabs were hovering over *E.coli* OP50 lawns without contacting them. (H-J) Thrashing assays of 3-day adult N2 wildtype (H) AWB-deficient (I) and ASH-deficient (J) worms in the presence (CL) or absence (M9) of worm corpse lysates. For (B-J): Solid circles represent individual biological replicates; hollow circles represent individual technical replicates (B-G) or individually assessed animals (H-J). Corresponding biological and technical replicates are color matched. Error bars represent standard deviation of the mean. Indicated P values were calculated using 1-way ANOVA tests with multiple comparison with M9 control condition serving as reference (B-G), and unpaired two-sided t tests (H-J). ns=p>0.05 (not significant); *p=<0.05; **p=<0.01; ***p=<0.001.

**Figure S4. Olfactory neurons AWB and ASH are required for dead perception-induced behavioral and physiological changes.** (A-L) Four-option choice assays testing behavior of 1-day old chooser animals upon exposure to worm corpse lysates. In each experiment, three *E.coli* OP50 lawns were supplemented with 10μg worm lysate whereas the fourth *E.coli* OP50 lawn was supplemented with M9 and served as control. Tested choosers were N2 wildtype (A), *che-2(e1033)* (B), and animals deficient in AWA & AWC (C), AWA (D), ADF (E), AQR, PQR, URX (F), ASK (G), AWC (H), ASI (I), ASJ (J), ASG/BAG neurons (K), or ASE (L). For (A-L): Solid circles represent individual biological replicates; hollow circles represent individual technical replicates. Corresponding biological and technical replicates are color matched. Error bars represent standard deviation of the mean. Indicated P values were calculated using 1-way ANOVA tests with multiple comparison with M9 control condition serving as reference. ns=p>0.05 (not significant); *p=<0.05; **p=<0.01; ***p=<0.001.

**Figure S5. Death perception involves glutaminergic signaling, GPCR activity, and cGMP-gated channels.** (A-R) Four-option choice assays testing behavior of 1-day old chooser animals upon exposure to worm corpse lysates. In each experiment, three *E.coli* OP50 lawns were supplemented with 10μg worm lysate whereas the fourth *E.coli* OP50 lawn was supplemented with M9 and served as control. Tested choosers were N2 wildtype (A), and animals deficient in *unc-25* (B), *unc-47* (C), *tbh-1* (D), *tdc-1* (E), *ptps-1* (F), *tph-1* (G), *bas-1* (H), *dop-3* (I), *cat-4* (J), *eat-4* (K), *osm-9* (L), *ocr-2* (M), *nlg-1* (N), *nmr-1* (O), *odr-1* (P), AIA neurons (Q), AIB neurons (R), AIY neurons (S), AIA & AIB neurons (T), or AIY & AIB neurons (U). For (A-U): Solid circles represent individual biological replicates; hollow circles represent individual technical replicates. Corresponding biological and technical replicates are color matched. Error bars represent standard deviation of the mean. Indicated P values were calculated using 1-way ANOVA tests with multiple comparison with M9 control condition serving as reference. ns=p>0.05 (not significant); *p=<0.05; **p=<0.01; ***p=<0.001.

**Figure S6. Characterization of death cue.** (A-J) Four-option choice assays in which 1-day old adult choosers were exposed to indicated corpse lysates. In (A), worms were exposed to chemically inactivated *C. briggsae* (isolate 1983). In (B), *E.coli* OP50 lawns were supplemented with indicated quantities of *Schmidtea mediterranea* (planaria) lysates. For (C), selected OP50 lawns were supplemented with conditioned M9 in which worms were previously cultured for 60 minutes, or CL1 corpse lysates. In (D), *E.coli* OP50 lawns were supplemented with M9 buffer, or lysates of worms inactivated with heat (65°C, 1hour) (CL5), NaN_3_ (CL1), or homogenized without prior inactivation (CL2). For (E-G), utilized worm corpse lysates were freeze-thawed five times (E), boiled (100°C, 30min) (F), or kept at room temperature for 3 weeks (G) prior to testing them in the depicted assays. In (H-I), worm corpse lysates were treated with DNAse (H) or proteinase K (I), respectively, prior to testing them in four-option choice assays. In (J), *E.coli* OP50 lawns were spiked with M9 buffer, CL2, Methanol/water fraction or methanol fraction from polarity-based CL1 fractionations. Fractionated metabolites were freeze-dried and resuspended in M9 prior to testing. (K-L). NMR spectra of metabolite identification assays using complete CL1 lysate (K) or the methanol fraction of CL1 lysate (L). For (A-J): Solid circles represent individual biological replicates; hollow circles represent individual technical replicates. Corresponding biological and technical replicates are color matched. Error bars represent standard deviation of the mean. Indicated P values were calculated using 1-way ANOVA tests with multiple comparison with M9 control condition serving as reference. ns=p>0.05 (not significant); *p=<0.05; **p=<0.01; ***p=<0.001.

**Figure S7. Testing of metabolites as potential death cues.** (A-F) Four-option choice assays in which 1-day old adult choosers were exposed to indicated conditions. In (A), worms were exposed to 10μl M9, 10μl CL5, 10μl of 100 mM lactate, and 10μl CL5 supplemented with lactate to a final concentration of 100mM. For (B), worms were exposed to 10μl M9, 10μl CL5, 10μl of 100 mM Nicotinamide adenine dinucleotide (NAD), and 10μl CL5 supplemented with NAD to a final concentration of 100mM. (C-F): worms were exposed to 10μl M9, 10μl CL1, 10μl of 100 mM cysteine (Cys) and alanine (Ala) (C), 10μl of 100 mM histidine (His) and Arginine (Arg) (D), 10μl of 100 mM glycine (Gly) and glucose (Gluc.) (E) or 10μl of 100 mM cysteine (Cys) and AMP (F). For (A-F): Solid circles represent individual biological replicates; hollow circles represent individual technical replicates. Corresponding biological and technical replicates are color matched. Error bars represent standard deviation of the mean. Indicated P values were calculated using 1-way ANOVA tests with multiple comparison with M9 control condition serving as reference. ns=p>0.05 (not significant); *p=<0.05; **p=<0.01.

