## Supplementary figures and images for "*C. elegans* behavior, fitness, and lifespan, are modulated by AWB/ASH-dependent death perception"

### Supplementary Figure S1

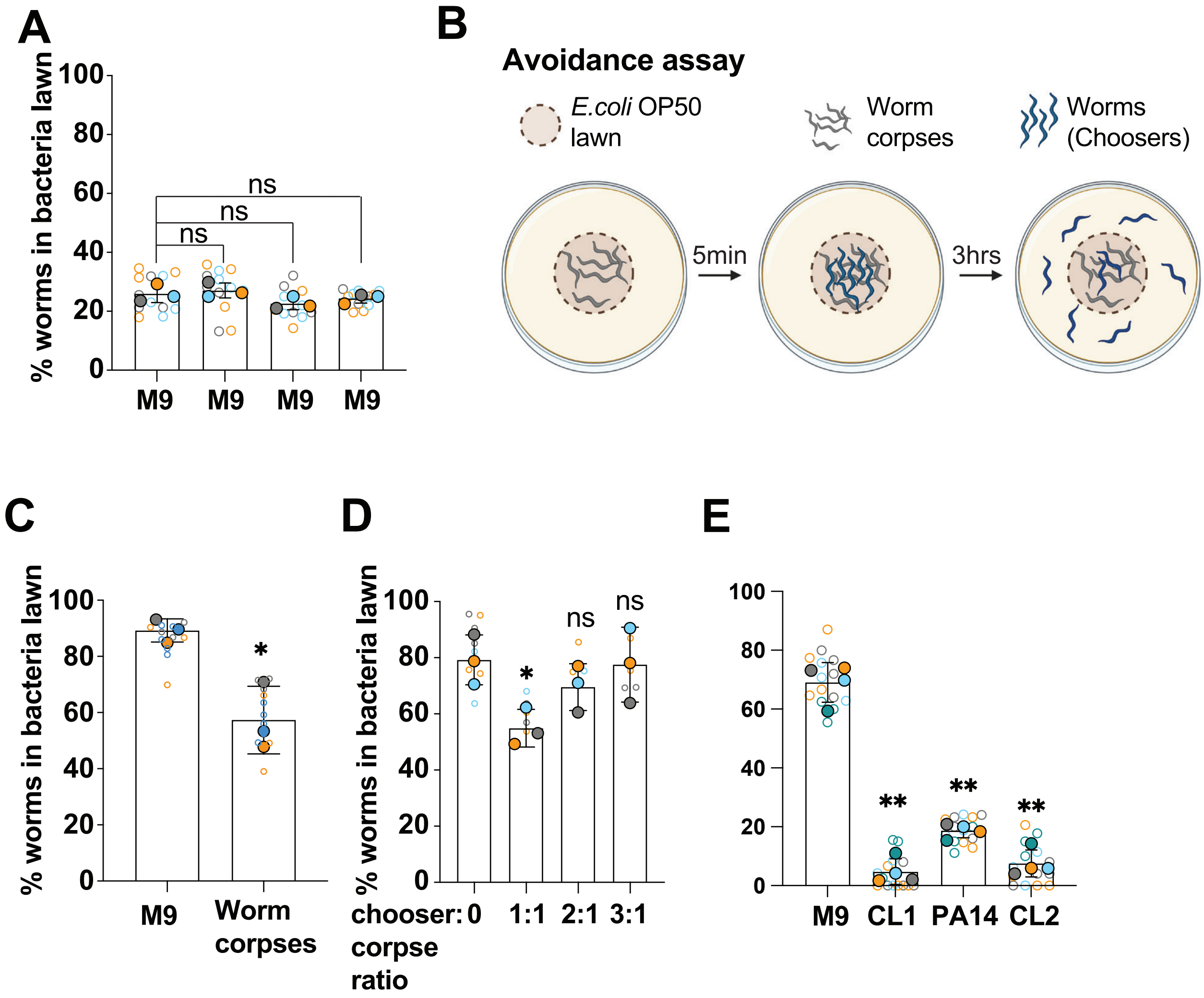

### Supplementary Figure S2

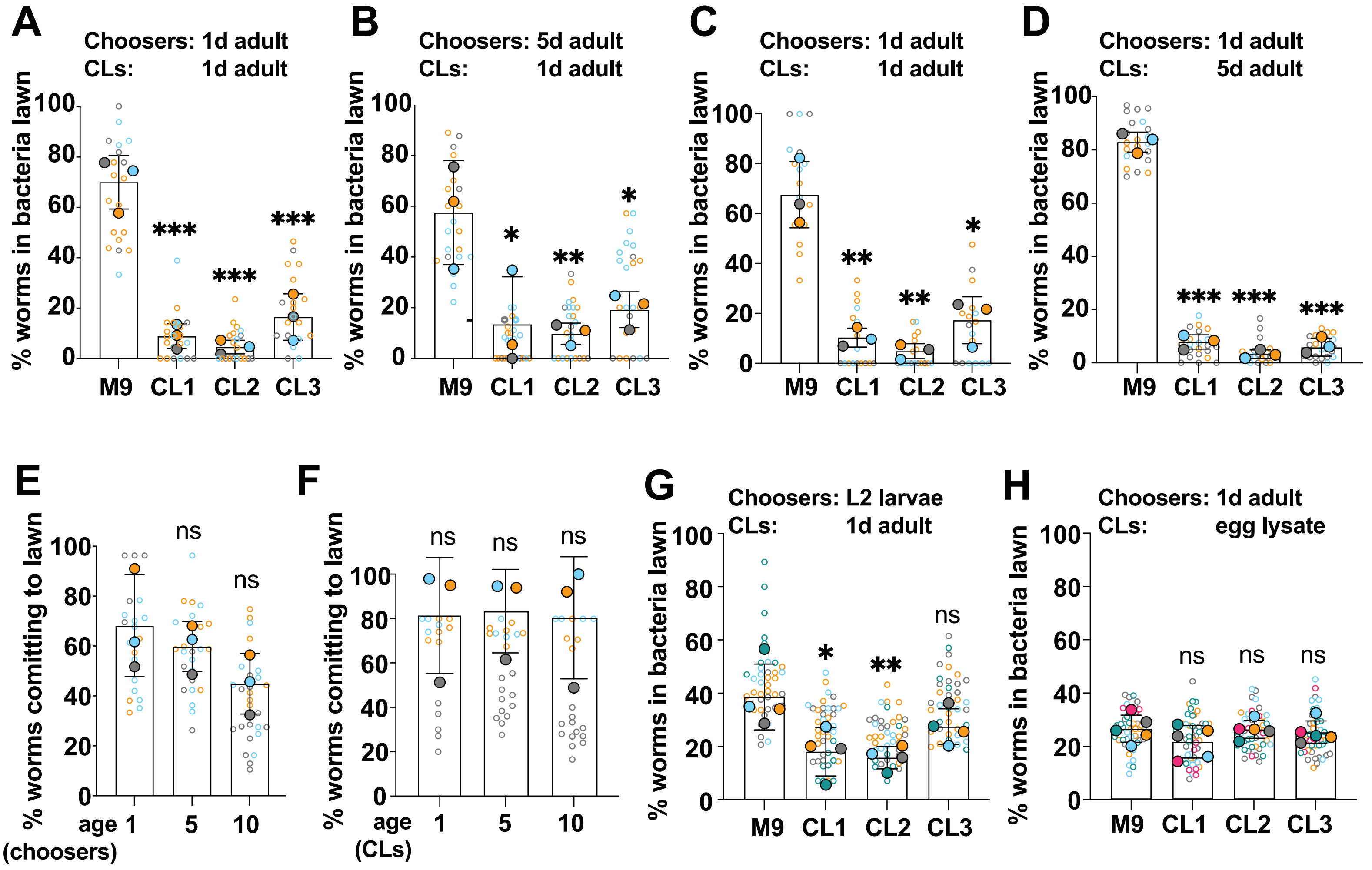

### Supplementary Figure S3

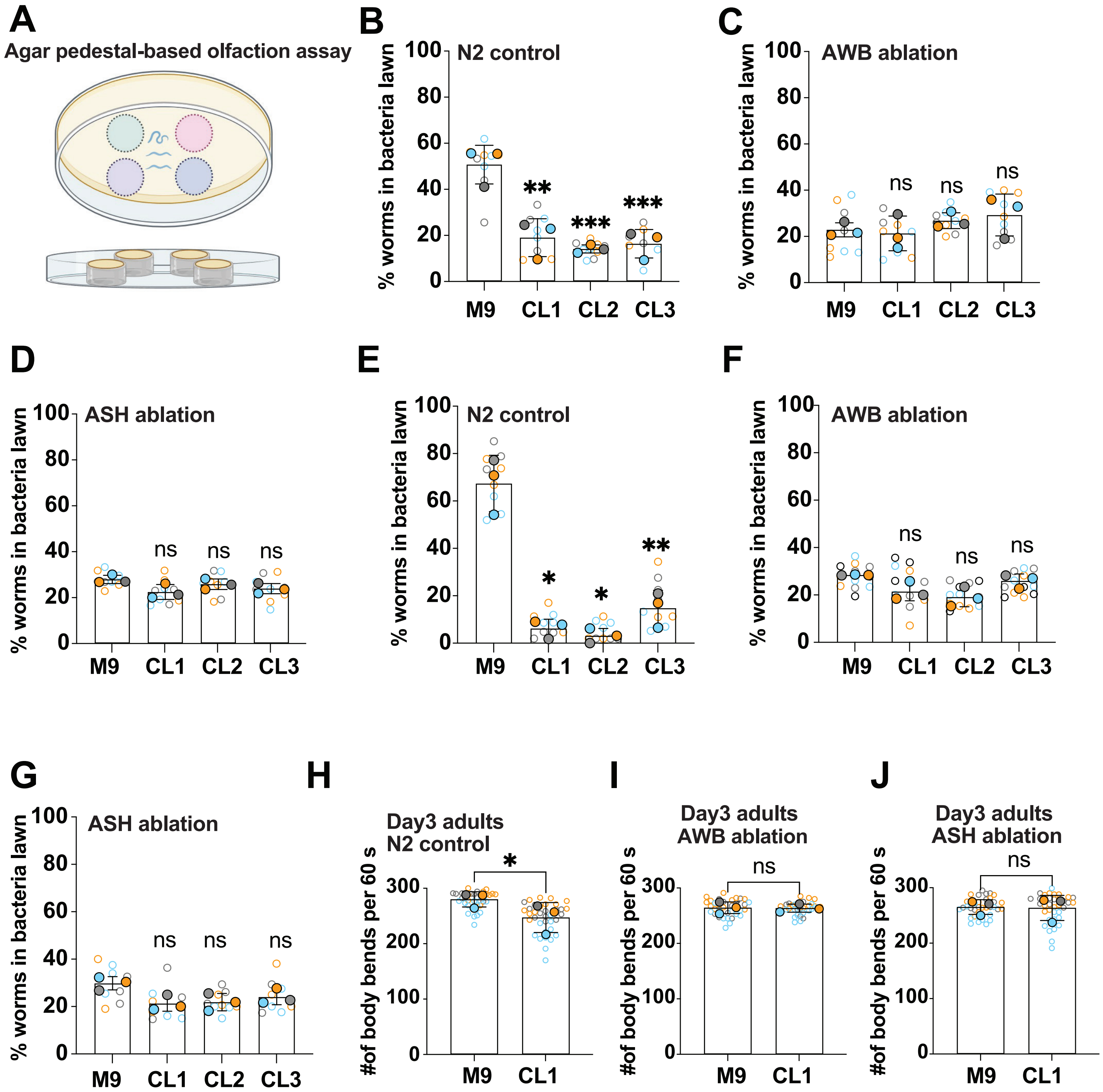

### Supplementary Figure S4

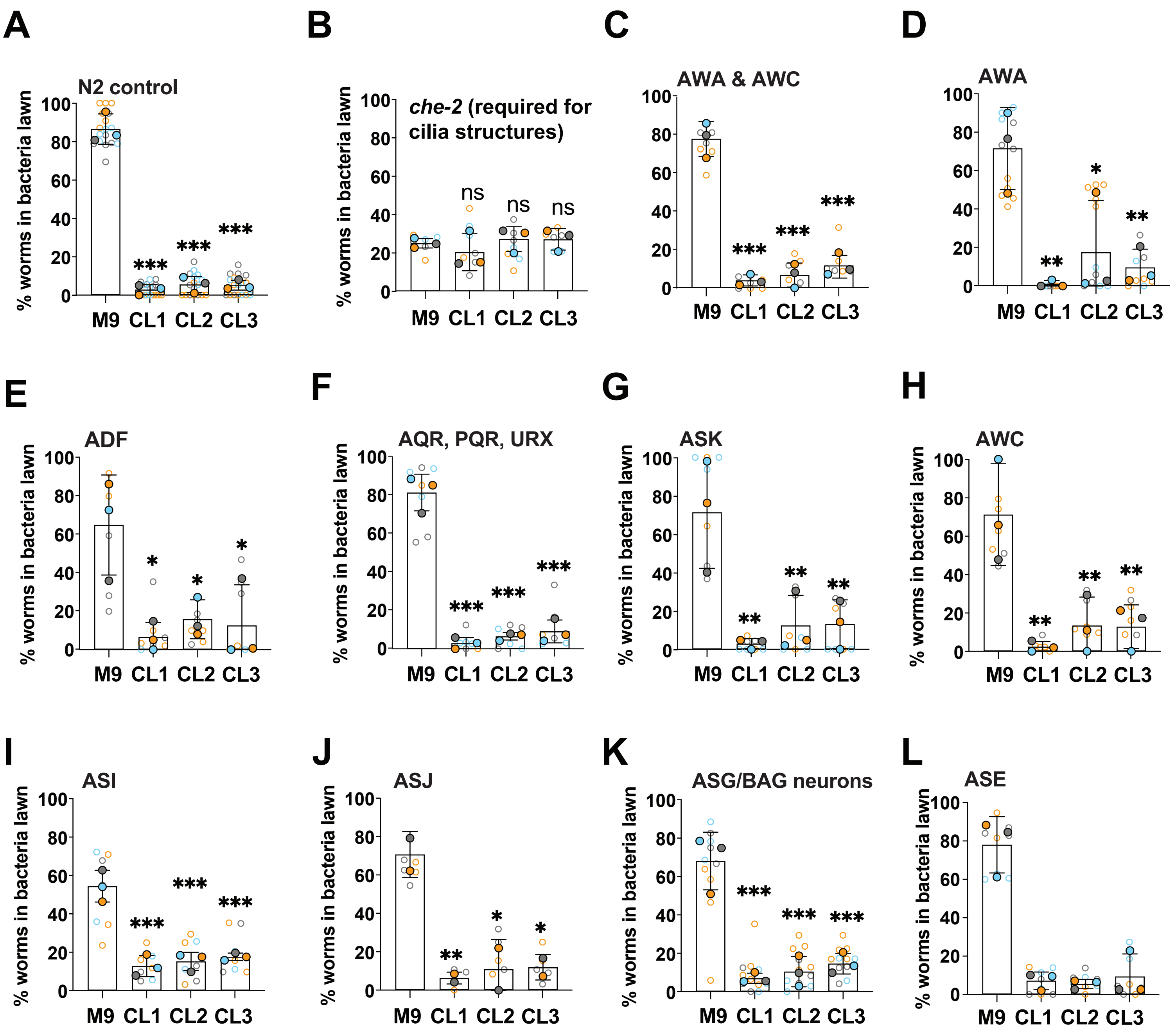

### Supplementary Figure S5

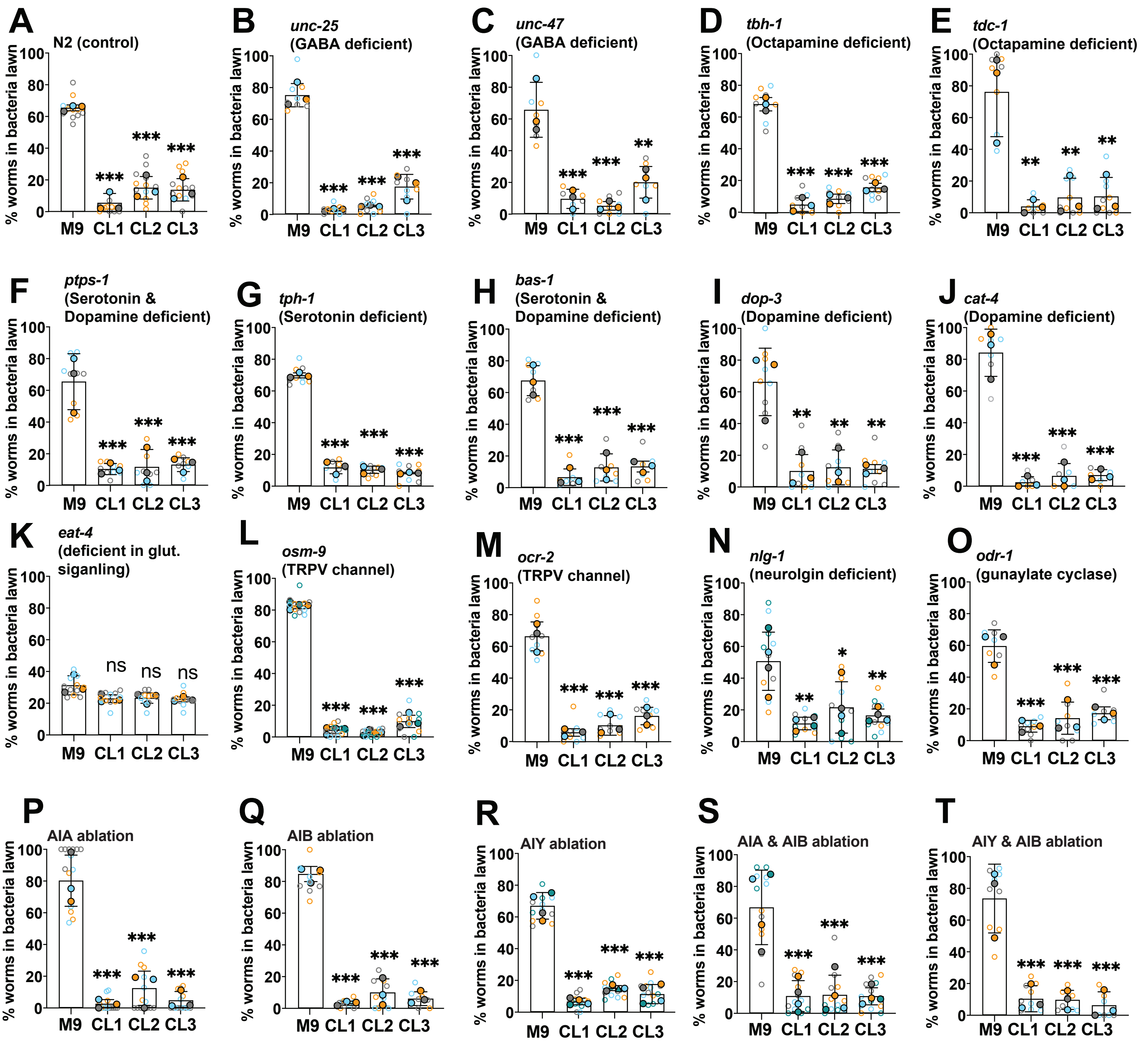

### Supplementary Figure S6

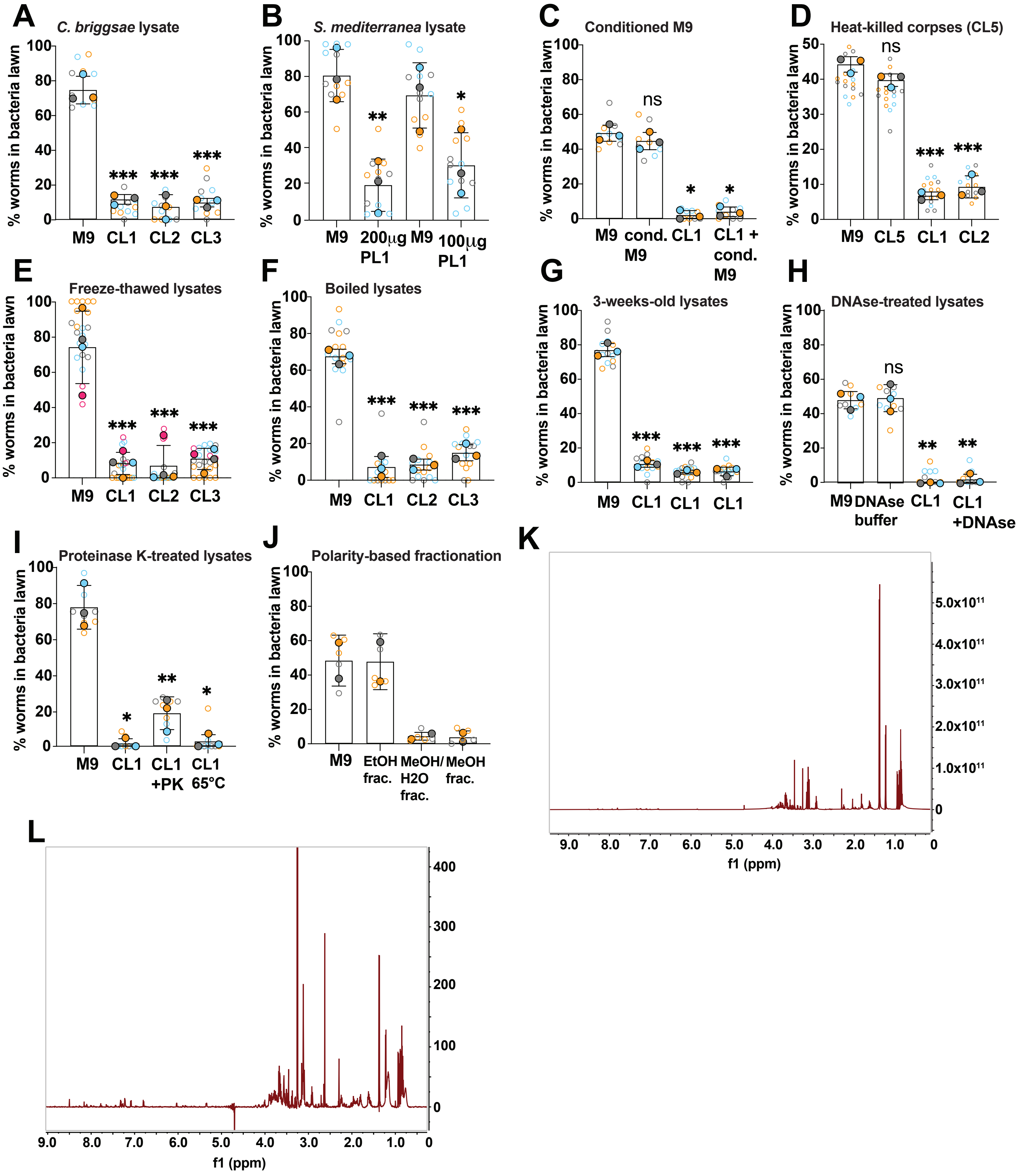

### Supplementary Figure S7

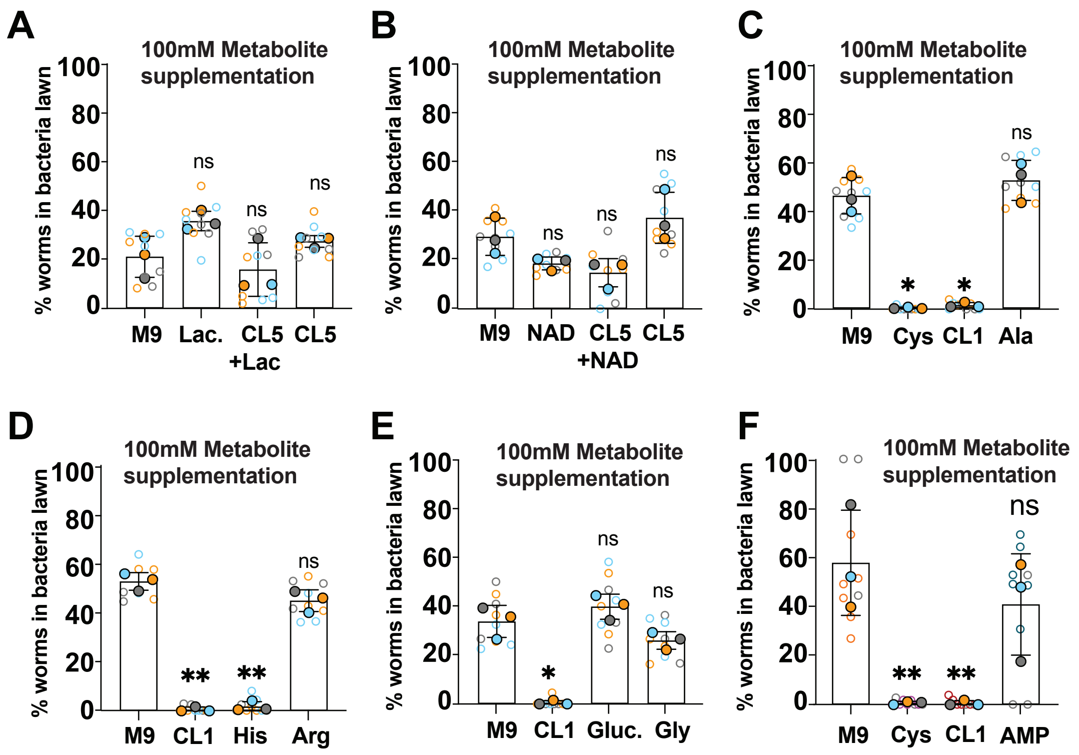
