## Supplemental Table S1 for "*C. elegans* behavior, fitness, and lifespan, are modulated by AWB/ASH-dependent death perception"

**Unfractionated CL1 lysate**

| Metabolite | Diagnostic peak (ppm) |
|---|---|
| S-Adenosylhomocysteine | 8.4, 8.3, 6.1, 4.9, 4.4, 4.3, 3.8, 3.1, 3.0, 2.7, 2.1 |
| AMP | 8.6, 8.2, 6.1, 4.8, 4.5, 4.4, 4.0 |
| Histidine | 7.9, 7.1, 4.0, 3.2, 3.1 |
| Methylhistidine | 8.0, 7.1, 3.9, 3.3, 3.2 |
| Tryosine | 7.2, 6.9, 3.9, 3.2, 3.0 |
| Phenylalanine | 7.4, 7.3, 4.0, 3.3, 3.1 |
| Uracil | 7.5, 5.8 |
| Glutamate | 3.8, 2.4, 2.3, 2.1, 2.0 |
| Aspartate | 3.9, 2.8, 2.7 |
| Cysteine | 4.0, 3.1, 3.0 |
| Glycine | 3.6 |
| Methanol | 3.4 |
| Agmatine | 7.2, 3.2, 3.0, 1.7 |
| Glucose | 5.2, 4.6, 3.9, 3.8, 3.7, 3.5, 3.4, 3.2 |
| Alanine | 3.8, 1.5 |
| Lactate | 4.1, 1.3 |
| Valine | 3.6, 2.3, 1.0 |
| leucine | 3.7, 1.7, 1.0, 0.9 |
| Succinate | 2.4 |
| Arginine | 7.2, 6.7, 3.8, 3.2, 1.9, 1.7, 1.6 |
| Isoleucine | 3.7, 2.0, 1.5, 1.2, 1.0, 0.9 |
