## Supplemental Table S3 for "*C. elegans* behavior, fitness, and lifespan, are modulated by AWB/ASH-dependent death perception"

**Table S3. Mass spectrometry-based metabolite identification. NaN3-0910 refers to unprocessed CL1-type lysate. NaN3-MeOH2 refers to the MeOH fraction type lysate. Orange highlighted metabolites were found in both complete corpse lysate and the active MeOH lysate fraction.**

| # | shared name | Compound_Name | Adduct | precursor mass |
|---|---|---|---|---|
| 1 | 18422 | [2-hydroxy-3-[hydroxy-[2,3,4,5,6-pentahydroxycyclohexyl]oxyphosphoryl]oxypropyl] octadecanoate | M-H | 599.3199 |
| 2 | 1494 | Adenosine monophosphate (AMP) | M-H | 346.0531 |
| 3 | 712 | Lyso PC (16:1) | M+H | 494.3221 |
| 4 | 9699 | Lyso PC (20:5) | M+H | 542.3243 |
| 5 | 11370 | Massbank:LQB00600 PE 38:6 | M-H | 762.5001 |
| 6 | 2922 | Massbank:PR309158 LPC 18:2 | M+HCOO | 564.3269 |
| 7 | 18346 | Massbank:PR309160 LPC 18:1 | M+HCOO | 566.3398 |
| 8 | 32 | Massbank:PR310844 LPC 18:2 | M+H | 520.3393 |
| 9 | 9759 | Monomyristin | M+H-H2O | 285.2425 |
| 10 | 9846 | PC-DAG (16:0/18:5) | M+H | 752.5122 |
| 11 | 9701 | Spectral Match to 1-(9Z-Octadecenoyl)-sn-glycero-3-phosphocholine from NIST14 | M+H | 522.3516 |
| 12 | 1486 | Spectral Match to 1-(9Z-Octadecenoyl)-sn-glycero-3-phosphoethanolamine from NIST14 | M-H | 478.2889 |
| 13 | 9743 | Spectral Match to 1-Heptadecanoyl-sn-glycero-3-phosphocholine from NIST14 | M+H | 510.3498 |
| 14 | 9703 | Spectral Match to 1-Hexadecanoyl-sn-glycerol from NIST14 | M+H | 331.2847 |
| 15 | 319 | Spectral Match to 1-O-Hexadecyl-2-O-(2E-butenoyl)-sn-glyceryl-3-phosphocholine from NIST14 | M+H | 550.3855 |
| 16 | 18397 | Spectral Match to 1-Palmitoyl-2-hydroxy-sn-glycero-3-phosphoethanolamine from NIST14 | M-H | 452.2726 |
| 17 | 9822 | Spectral Match to 1-Pentadecanoyl-sn-glycero-3-phosphocholine from NIST14 | M+H | 482.324 |
| 18 | 90 | Spectral Match to 1-Stearoyl-2-hydroxy-sn-glycero-3-phosphocholine from NIST14 | M+H | 524.3702 |
| 19 | 5370 | Spectral Match to 1-Stearoyl-2-hydroxy-sn-glycero-3-phosphoethanolamine from NIST14 | M-H | 480.3052 |
| 20 | 9793 | Spectral Match to 1,2-Di-(9Z,12Z,15Z-octadecatrienoyl)-sn-glycero-3-phosphocholine from NIST14 | M+H | 778.5344 |
| 21 | 10002 | Spectral Match to 1,2-Diarachidonoyl-sn-glycero-3-phosphocholine from NIST14 | M+H | 830.5784 |
| 22 | 24 | Spectral Match to 13-Docosenamide, (Z)- from NIST14 | M+H | 338.3415 |
| 23 | 9913 | Spectral Match to 14,15-EE-(8Z)-E from NIST14 | M+H-H2O | 307.2581 |
| 24 | 9774 | Spectral Match to 8S-Hydroxy-9E,11Z,14Z-eicosatrienoic acid from NIST14 | M+H-H2O | 305.2463 |
| 25 | 9729 | Spectral Match to Dimethyldioctadecylammonium cation from NIST14 | Cat | 550.6216 |
| 26 | 9936 | Spectral Match to Docosahexaenoyl PAF C-16 from NIST14 | M+H | 792.5496 |

| | | | | |
|---|---|---|---|---|
| 27 | 9780 | Spectral Match to Glycerol 1-myristate from NIST14 | M+H | 303.2478 |
| 28 | 9702 | Spectral Match to Glycerol 1-stearate from NIST14 | M+H | 359.3104 |
| 29 | 10421 | Spectral Match to His-Trp from NIST14 | M+H | 342.1557 |
| 30 | 10466 | Spectral Match to Ile-Gly-Ile from NIST14 | M+H | 302.202 |
| 31 | 10489 | Spectral Match to Ile-Pro-Ile from NIST14 | M+H | 342.235 |
| 32 | 9795 | Spectral Match to Leu-Phe from NIST14 | M+H | 279.1692 |
| 33 | 10500 | Spectral Match to Leu-Trp from NIST14 | M+H | 318.176 |
| 34 | 9828 | Spectral Match to Monoelaidin from NIST14 | M+H-H2O | 339.2842 |
| 35 | 10042 | Spectral Match to Stearidonic acid from NIST14 | M+H | 277.2121 |
| 36 | 10109 | Spectral Match to Tyr-Phe from NIST14 | M+H | 329.1458 |
| 37 | 9948 | SPERMIDINE | M+H | 146.1644 |
| 38 | 5070 | TOP19 Psoriasis feature - Unknown FeatureID=3668 | M+H | 466.3263 |

**obtained by polarity based fractionation of CL1-**

| RTMean | Sample Distribution |
|---|---|
| 5.5963 | NaN3-0910 |
| 0.4835 | NaN3-MeOH2,NaN3-0910 |
| 4.9287 | NaN3-0910 |
| 4.9287 | NaN3-0910 |
| 7.8659 | NaN3-0910 |
| 5.084 | NaN3-MeOH2,NaN3-0910 |
| 5.2915 | NaN3-0910 |
| 4.9563 | NaN3-MeOH2 |
| 5.5655 | NaN3-0910 |
| 7.9075 | NaN3-0910 |
| 5.2887 | NaN3-0910 |
| 5.1394 | NaN3-MeOH2,NaN3-0910 |
| 5.344 | NaN3-0910 |
| 6.0085 | NaN3-0910 |
| 5.6349 | NaN3-MeOH2,NaN3-0910 |
| 5.0698 | NaN3-0910 |
| 4.9564 | NaN3-0910 |
| 5.4827 | NaN3-MeOH2 |
| 5.4303 | NaN3-0910 |
| 7.836 | NaN3-0910 |
| 8.7523 | NaN3-0910 |
| 6.868 | NaN3-MeOH2 |
| 6.3962 | NaN3-0910 |
| 6.2024 | NaN3-0910 |
| 8.2236 | NaN3-0910 |
| 9.1373 | NaN3-0910 |

| | |
|---|---|
| 5.5655 | NaN3-0910 |
| 6.5069 | NaN3-0910 |
| 0.8984 | NaN3-0910 |
| 2.5199 | NaN3-0910 |
| 2.6727 | NaN3-0910 |
| 2.6306 | NaN3-0910 |
| 2.7691 | NaN3-0910 |
| 6.2024 | NaN3-0910 |
| 5.787 | NaN3-0910 |
| 2.4922 | NaN3-0910 |
| 0.3447 | NaN3-0910 |
| 5.5397 | NaN3-0910 |
