## Supplemental Table S4 for "*C. elegans* behavior, fitness, and lifespan, are modulated by AWB/ASH-dependent death perception"

| Strains | Genotype | Description | Figure |
|---|---|---|---|
| N2 Bristol | Wild-type | All experiments | All experiments |
| BC1977 | *C. briggsae* | Cbr-unc(s1275) | Figure 6; Panel C |
| BC1983 | *C. briggsae* | Cbr-dpy(s1281) | Supplementary Figure 6; Panel A |
| BZ873 | *dop-3(ok295) X* | Dopamine receptor knockout | Supplementary Figure 5; Panel I |
| CB1033 | *che-2(e1033) X* | Chemotaxis abnormal. Amphid defect-EM | Supplementary Figure 4; Panel B |
| CB1124 | *che-3(e1124) I.* | Chemotaxis abnormal | Figure 4; Panel A |
| CX11697 | *kyls536;kyls538* | Death of ASG as well as BAG neurons | Supplementary Figure 4; Panel K |
| CX2065 | *odr-1(n1936) X* | Defective chemotaxis to some volatile odorants: benzaldehyde, 2-butanone, isoamyl alcohol | Supplementary Figure 5; Panel P |
| CX4544 | *ocr-2(ak47) IV.* | Chemosensory, mechanosensory, and osmosensory defects | Supplementary Figure 5; Panel M |
| DR47 | *daf-11(m47) V.* | Chemotaxis defective (Na+) | Figure 5; Panel H |
| FG540 | *N2; udEx428[srh-124p::ced-3(p15), srh-142p::ced-3(p17), srh-142p::gfp, elt-2p::gfp]* | Ablation of ADF | Supplementary Figure 4, Panel E |
| FK100 | *tax-2(ks10) I* | Contributes to intracellular cGMP-activated cation channel activity | Figure 5; Panel F |
| FK103 | *tax-4(ks28) III* | Contributes to intracellular cGMP-activated cation channel activity | Figure 5; Panel G |
| FK223 | *egl-4(ks60) IV* | Larger body size, longer lifespan, egg-laying defect | Figure 5; Panel E |
| IK130 | pkc-1(nj3) V. | Defective chemotaxis to AWA- and AWC-sensed odorants | Figure 4,Panel C |
| IK600 | *eat-4(nj2) III* | Involved in neurotransmitter transmembrane transporter activity | Figure 5; Panel D |
| IK602 | *eat-4(nj6) III.* | Involved in neurotransmitter transmembrane transporter activity | Supplementary Figure 5; Panel K |
| JN1713 | *peIs1713 [sra-6p::mCasp-1 + unc-122p::mCherry].* | ASH neurons are eliminated | Figure 4; Panels E & I, Supplementary Figure 3; Panels D,G, & J |
| JN1715 | *peIs1715* | AWB neurons are eliminated | Figure 4; Panels D,F & H, Supplementary Figure 3; Panel C,F,& I |

| | | | |
|---|---|---|---|
| JN578 | *peIs578* | AIB neurons are ablated by specific expression of caspase | Supplementary Figure 5; Panel R |
| JN579 | *peIs579* | AIY neurons are ablated by specific expression of caspase | Supplementary Figure 5; Panel S |
| JN580 | *peIs580* | AIA neurons are ablated by specific expression of caspase | Supplementary Figure 5; Panel Q |
| JN604 | *pels578[npr-9p::casp1 npr-9::venus unc-122 p::mCherry];pels579 [tt-3::casp1 ttx-3p::venus lin-44::gfp]* | Genetic ablation of AIY/AIB | Supplementary Figure 5; Panel U |
| JN605 | *pels578[npr-9p::casp1 npr-9::venus unc-122 p::mCherry];pels580 [ins-1(short)p::casp1 ins-1 (short)::gfp unc-122p::gfp]* | Genetic ablation of AIB/AIA | Supplementary Figure 5; Panel T |
| KC435 | *C. remanei* | Male-female strain | Figure 6; Panel B |
| LC33 | *bas-1(tm351) III* | Serotonin-deficient by anti-serotonin straining | Supplementary Figure 5; Panel H |
| LC80 | *ptps-1(tm1984) I.* | Serotonin- and dopamine-deficient, tetrahydrobiopterin-deficient, general chemical hypersensitivity | Supplementary Figure 5;Panel F |
| LC81 | *cat-4(tm773) V.* | Serotonin and dopamine-deficient | Supplementary Figure 5;Panel J |
| MT13113 | *tdc-1(n3419) II* | n3419 is a deletion in L-aromatic amino acid decarboxylase with homology to histadine decarboxylase. | Supplementary Figure 5; Panel E |
| MT14984 | *tph-1(n4622) II* | Egl. Reduced pharyngeal pumping | Supplementary Figure 5; Panel G |
| MT6201 | *unc-47(n2409) III.* | Involved in GABAergic synaptic transmission | Supplementary Figure 5; Panel C |
| MT6490 | *unc-25(n2569) III* | Involved in GABAergic synaptic transmission | Supplementary Figure 5; Panel B |
| MT9455 | *tbh-1(n3247) X* | n3247 is a 791 bp deletion which results in a truncated TBH-1 protein. Hypersensitive to 5-HT. Reduced locomotion rate. | Supplementary Figure 5; Panel D |
| OH8585 | *otIs4 [gcy-7::GFP]. otEx3822 [ceh-36::CZ-caspase3(p17) + gcy-7::caspase3(p12)-NZ + myo-3::mCherry]* | otIs4 [gcy-7::GFP]. otEx3822 [ceh-36::CZ-caspase3(p17) + gcy-7::caspase3(p12)-NZ + myo-3::mCherry] | supplementary Figure 4, Panel L |
| PS6025 | *qrIs2* | qrIs2 [sra-9::mCasp1]. Caspase expression in ASK neuron | Supplementary Figure 4; Panel G |

| | | | |
|---|---|---|---|
| PY7502 | *oyIs85* | AWC ablated in this strain. Defective thermotaxis | Supplementary Figure 4; Panel H |
| PY7505 | *oyIs84.* | ASI is ablated in this strain | Supplementary Figure 4; Panel I |
| VC1262 | *osm-9(ok1677) IV* | Enables calcium channel activity and temperature-gated cation channel activity | Supplementary Figure 5; Panel L |
| VC228 | *nlg-1(ok259) X* | Involved in gamma-aminobutyric acid receptor clustering | Supplementary Figure 5; Panel N |
| VM487 | *nmr-1(ak4) II.* | Contributes to NMDA glutamate receptor activity | Supplementary Figure 5; Panel O |
| XA2262 | *gcy-33(ok232) V; gcy-31(ok296) X; qaIs2241* | qaIs2241 [gcy-36p::egl-1 + gcy-35p::GFP]; causes genetic ablation of AQR, PQR, and URX neurons | Supplementary Figure 4; Panel F |
| ZD1306 | *ptrx-1::ICE + pofm-I::GFP]* | Ablation of ASJ | Supplementary Figure 4, Panel J |
